# Reaction-diffusion condensation generates a regulatable landscape for self-organizing subcellular structures

**DOI:** 10.64898/2026.06.14.732133

**Authors:** Eden Chang, Zhejing Xu, Elliott W. Z. Weix, Scott M. Coyle

## Abstract

Cells self-organize complex internal architectures through an interplay between biochemical signaling and physical structure. Within this paradigm, reaction-diffusion signaling mechanisms create local concentration gradients, and protein condensates assemble in a concentration-dependent manner, providing a natural coupling for a landscape of composite active structures to emerge. Here, we systematically chart this landscape in human cells at scale using RIPPLE (Reaction-diffusion IDR Platform for Producing Living Emulsions), a synthetic modular system that tethers disordered, condensate-forming IDR sequences to a programmable, two-protein wave-generating reaction-diffusion (RD) circuit. Fusing IDRs to the RD Activator, its partner ATPase, or both, generates a vast array of self-organizing subcellular architectures—ranging from traveling condensation waves to oscillating droplet networks to persistent phase-separated macrostructures that pattern the entire cell. By systematically tuning the frequency and amplitude of the underlying RD waveforms, we reveal frequency-dependent transitions regulating dilute, condensed, and aggregated states specified by IDR sequence chemistry and miscibility. RIPPLE provides a platform for understanding and engineering protein condensation far from equilibrium, revealing how signaling and structure can cooperate in a minimal, two-protein system to generate a tunable spectrum of dynamic cellular structures.

## Introduction

Cells are exquisitely structured systems whose physical form supports functions essential for biology^1,2^. While cell shape provides an overall structural foundation to the cell, further subcellular organization partitions the cell into distinct zones, compartments, or architectures to support specific activities ^3–5^. For example, cells can polarize to establish a molecularly distinct front and back to support leading-edge motility, or to establish distinct apical and basal domains in the epithelium to organize tissue barriers and nutrient uptake ^6–10^. Likewise, many unicellular protists and parasites create geometrically ordered arrays of motile cilia to perform niche-specific tasks, and similar patterns are established in human multiciliated cells to support efficient mucociliary transport ^5,11–14^. How cells generate, organize, and regulate this diversity of elaborate subcellular structures remains a fundamental question in biology^15^.

An important conceptual framework that has emerged is that patterns of spatiotemporal signaling at the subcellular level can organize target architectures by locally controlling access to factors that regulate the assembly or disassembly of downstream structures ^16–18^. For example, intrinsic cell polarity depends on self-organizing signaling mechanisms to create a localized pool of active small GTPases that regulate actin polymerization^19–21^. The patterns these signaling systems create arise through a combination of reaction and diffusion of their molecular components. Localized GTPase activation (through positive feedback) and global inhibition can provide a winner-take-all control logic for polarization^20,22^. Alternatively, traveling waves of Rho GTPase activity arise through a Turing-type reaction-diffusion mechanism, in which non-linear reaction kinetics and differences in diffusion rates between cytosolic and membrane-bound states lead to spontaneous protein pattern formation^23–25^.

However, the connection of signaling systems to downstream structural modules can fundamentally alter their activity. Rather than acting as passive receivers of upstream information, structural components can feed back into the signaling landscape by sequestering components, altering diffusion rates, or changing mechanical constraints^26–29^. Consequently, the coupling of chemical signaling with physical structure creates an environment poised for composite active structures to emerge^3,30^. Highly evolved architectures are often governed by dozens of overlapping regulatory inputs optimized for highly specific biological tasks, making it difficult to systematically isolate how different structure-signaling combinations alter emergent behavior^18,31,32^. In contrast, synthetic biology provides a bottom-up alternative where we can construct and characterize orthogonal systems built from modular signaling patterns and structural elements to map the landscape of outputs directly in living cells at scale ^18,33–39^.

Biomolecular condensates provide an ideal, minimal class of micron-scale structures to explore this way synthetically, because assembly and disassembly are directly controlled by local effective concentration ^40–44^. In living systems, these condensates are frequently coupled to energy-consuming signaling networks that drive continuous molecular exchange and spatial reorganization—ranging from ATP-dependent chromatin remodeling to T-cell synapse formation^45–47^. Yet, while numerous studies have characterized condensate behavior at thermodynamic equilibrium, how they behave and the structures they generate when driven by spatiotemporally shifting concentration gradients remain poorly understood^48,49^.

To access and systematically explore this non-equilibrium regime at scale, we developed **RIPPLE** (Figure 1A): a minimal, fully synthetic system operating in human cells that couples a tunable reaction-diffusion engine to protein condensates. To achieve this, we leverage our recent adaptation of the bacterial MinDE divisome positioning system as an orthogonal, programmable reaction-diffusion (RD) system in human cells^37,50,51^. Expression of the MinD ATPase and its Activator, MinE, generates fast protein waves, oscillations, and patterns on membranes programmed by component levels^52–56^. By coupling intrinsically disordered region (IDR) condensate-forming sequences to the MinDE machinery, RIPPLE establishes a synthetic platform that directly links phase separation to an ATP-driven, pattern-forming RD network.

**Figure 1:**
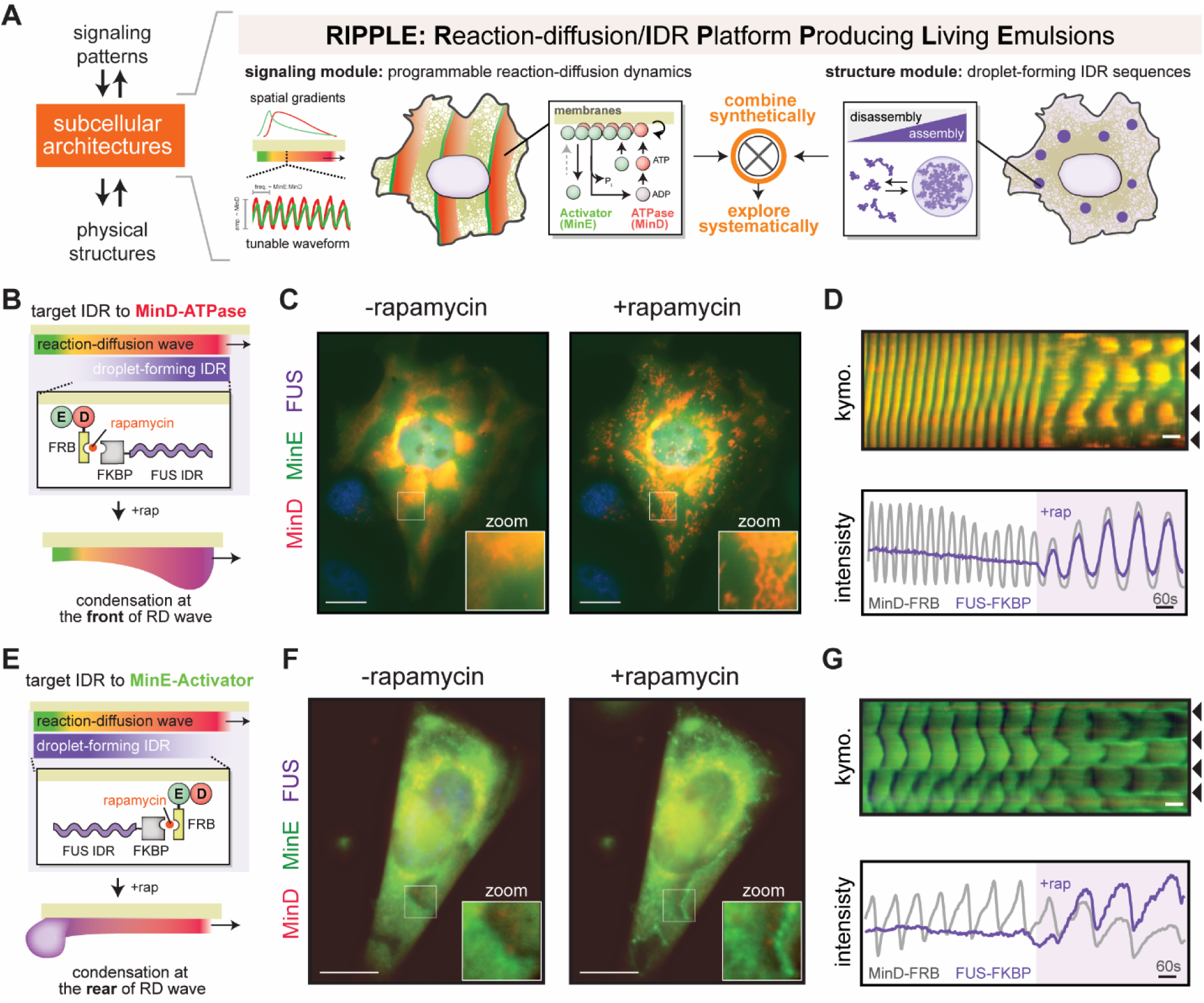
RIPPLE couples reaction–diffusion dynamics to biomolecular condensation. (A) Conceptual framework of the **R**eaction–diffusion/**I**DR **P**latform **P**roducing **L**iving **E**mulsion (RIPPLE). RIPPLE synthetically couples a signaling module, represented by reaction–diffusion (RD) dynamics, with a structural module composed of droplet-forming intrinsically disordered regions (IDRs) to co-regulate subcellular organization. The platform uses the minimal two-protein MinDE system, in which the ATPase MinD and its activator MinE generate programmable RD dynamics. (B–G) Rapamycin-inducible coupling of condensate-forming FUS-IDRs to the MinDE reaction–diffusion circuit. FUS-IDRs are recruited to MinD or MinE through FKBP–FRB heterodimerization, allowing RD waves to locally concentrate IDRs and trigger phase separation above a critical threshold. (B–D) Recruitment of FUS-IDRs to MinD. (B) Schematic of FUS-IDR-FKBP recruitment to FRB-MinD in the presence of wild-type MinE. (C) Representative fluorescence images before and after rapamycin induction. Condensates assemble at the leading edge of propagating waves following induction. Scale bar: 20 μm. (D) Corresponding kymograph and pixel-level time series analysis showing reduced oscillation frequency following induction. Kymograph scale bar: 60 s. (E–G) Recruitment of FUS-IDRs to MinE. (E) Schematic of FUS-IDR-FKBP recruitment to MinE-FRB in the presence of wild-type MinD. (F) Representative fluorescence images before and after rapamycin induction. Condensates assemble at the trailing edge of propagating waves following induction. Scale bar: 20 μm. (G) Corresponding kymograph and pixel-level time series analysis showing reduced oscillation frequency following induction. Kymograph scale bar: 60 s.

Using RIPPLE, we reveal a regulatable landscape of self-organizing structures defined by the interplay between reaction-diffusion (RD) and protein condensation, ranging from traveling condensation waves to oscillating droplet networks to persistent phase-separated patterns and macrostructures that span the cell. By systematically tuning the frequency and amplitude of the underlying RD waveforms, we define frequency-dependent transitions that regulate dilute, condensed, and aggregated states, as specified by IDR sequence chemistry and miscibility. The ease with which signaling and condensation cooperate in this minimal synthetic system suggests reaction-diffusion-condensation may be a more general paradigm governing the creation, diversification, and regulation of subcellular structure.

## Results

### Coupling droplet-forming IDR sequences to a reaction-diffusion system generates activity-driven condensation

Because reaction-diffusion systems create dynamic protein concentration gradients and protein condensate assembly is concentration-dependent, their coupling creates an opportunity for emergent structures and behaviors to emerge. To explore this coupling synthetically in human cells, we developed **RIPPLE**, a reaction-diffusion IDR platform for producing living protein emulsions. RIPPLE links droplet-forming IDR sequences to a programmable reaction-diffusion (RD) system based on the bacterial divisome system MinDE. Expression of the MinD ATPase and MinE activator generates synthetic protein waves whose frequency and amplitude are controlled by the expression levels of the components. These waves are driven by nucleotide-dependent assembly of the MinD ATPase at the wavefront and MinE-driven ATP hydrolysis and disassembly at the rear, creating micron-scale concentration gradients of both MinD and MinE. As such, coupling droplet-forming IDR sequences to the MinD or MinE proteins will impose time-varying concentration dynamics that push the IDR towards or away from condensation, while the reaction-diffusion kinetics of MinD and MinE might be altered within this condensed state (Fig. 1).

To explore this, we engineered an inducible RIPPLE platform that links droplet-forming intrinsically disordered regions (IDRs) to MinDE waves using FKBP/FRB recruitment with rapamycin^57^. This allows us to examine the appearance, structure, and dynamics of the reaction-diffusion wave both before and after the coupling of the IDR sequence. By fusing FRB to the MinD ATPase or the MinE activator, we can control whether the IDR is targeted to the assembling front or disassembling rear of the wave, respectively. As an initial IDR sequence, we selected FUS, a model droplet-forming sequence widely used to investigate protein condensation. In the absence of rapamycin, both FRB-MinD/E and MinD/E-FRB waves propagated normally in smooth spirals, and the FUS-IDR–FKBP-BFP remained diffusely distributed and did not participate in the oscillations (Fig. 1B-G; Movie S1-2). This establishes a baseline for comparison in which the condensation module is uncoupled from the reaction-diffusion system.

In transfected populations, we observed many cells in which recruitment of the FUS-IDR to MinD (front) led to rapid and dramatic changes in both wave appearance and dynamics (Fig. 1B-D; Movie S1). Within seconds, the previously smooth wavefronts of the uncoupled system became more irregular and separated into smaller, condensed clusters. Remarkably, these condensed waves continued to propagate throughout the cell, but with a more erratic and turbulent spatial structure (Fig. S1). This change in wave appearance correlated with a marked decrease in wave speed. Thus, in a D-FUS context, MinDE components continue to produce reaction-diffusion waves, but now appear to propagate through a more condensed medium that decreases both spatial coherence and wave speed.

Surprisingly, recruitment of FUS-IDR to MinE (rear) led to a completely different emergent behavior: droplets of MinE began to assemble in the wake of the reaction-diffusion wave while the front of the wave continued to propagate normally and smoothly with no apparent change in spatial wavelength (Fig. 1E-G; Fig. S1, Movie S2). In some cells, these MinE-FUS droplets appeared fixed in space, forming barriers that restricted or reflected wave propagation. In other cells, these droplets appeared to form at spiral wave centers, dynamically growing and dissolving in response to wave activity (Movie S2). As with D-IDR RIPPLE, changes in wave appearance correlated with a decrease in wave frequency—though in this case likely through sequestration of the E-IDR activator^54^.

Together, our inducible results establish that the RIPPLE system can effectively couple MinDE reaction-diffusion dynamics to condensate assembly and disassembly. The differences we see in behavior when targeting IDRs to the MinD front or the MinE rear reflect how condensation feeds back onto system dynamics, revealing distinct modes of interaction between phase separation and reaction–diffusion processes. This provides a starting point for systematic investigation into how different reaction-diffusion wave parameters and IDR sequences interact.

### Wave velocity acts as a low-pass filter that gates ATPase-IDR RIPPLE condensation

While our inducible recruitment experiments demonstrated that coupling FUS-IDR to the MinD ATPase wavefront can generate activity-driven condensation, this behavior varied across different cells in the transfected population. These differences could reflect variation in how the underlying frequency and amplitude of the reaction-diffusion wave modulate the local concentration dynamics of the IDR sequence. To systematically explore these effects, we took advantage of the fact that the MinDE waveform is set by expression levels, with the E/D ratio setting the frequency and the D levels setting the amplitude ^37,50,51^. This allows us to screen libraries of ATPase-IDR RIPPLE systems by fusing any droplet-forming IDR sequences directly to MinD (D-IDR) and scoring their emergent behaviors across a wide range of randomized expression levels (Fig. 2A).

**Figure 2:**
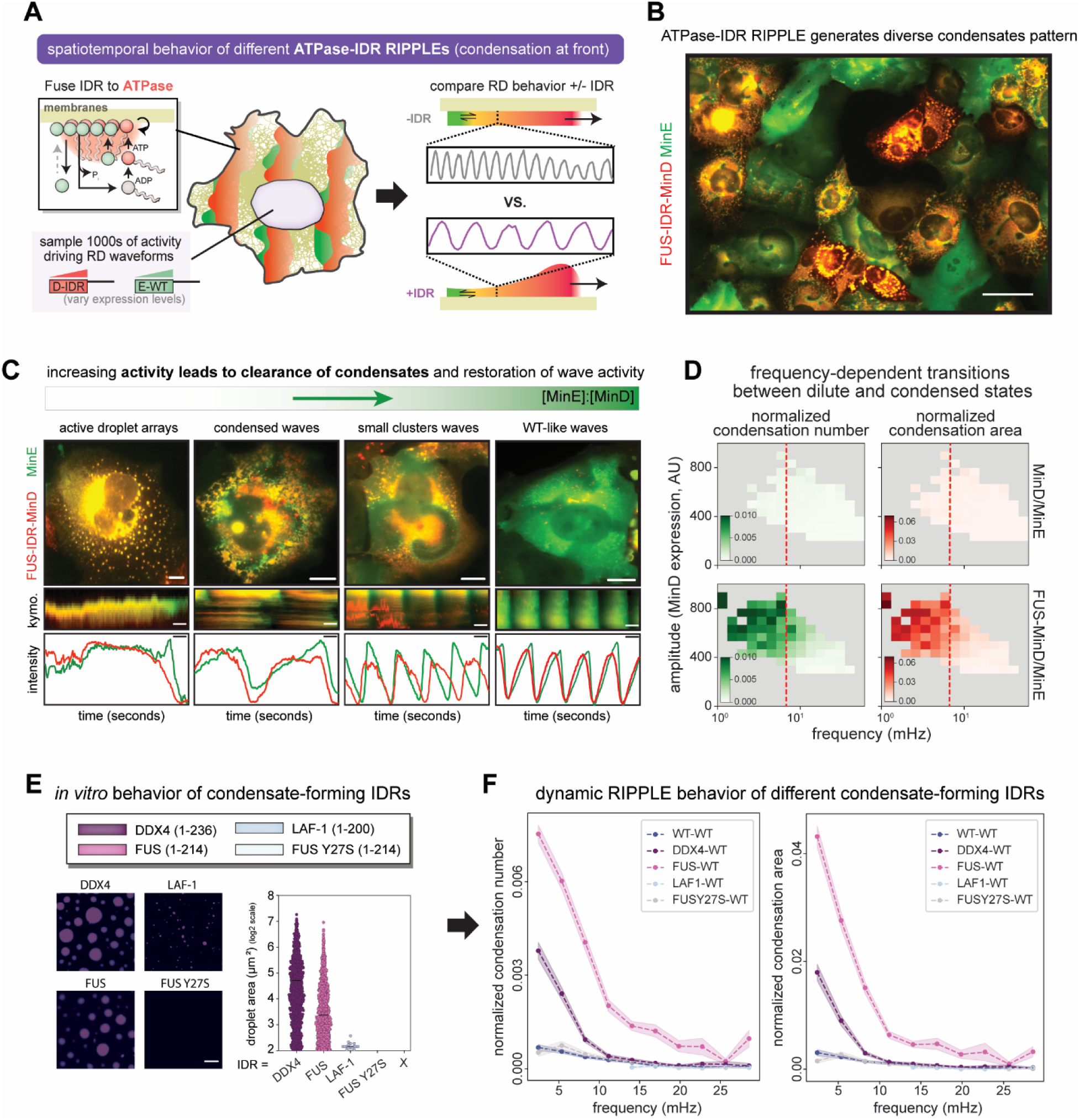
ATPase-IDR coupling reveals low-pass filtering of condensate assembly by reaction–diffusion dynamics. (A) Schematic of constitutive MinD-IDR RIPPLE design. IDR sequences are directly fused to MinD and co-expressed with wild-type MinE. (B) Representative fluorescence images of FUS-IDR-MinD expressing cells across a population, displaying diverse spatial organizations. Scale bar: 50 μm. (C) Representative single-cell images illustrating the transition of condensation behavior as a function of relative MinE-to-MinD (E/D) expression ratio. Increasing MinE levels drive a shift from stable, large condensates to dynamic propagating condensate waves, and ultimately to condensate-suppressed, WT-like oscillatory patterns. Scale bar: 10 μm. (D) Two-dimensional binned histogram of population condensation score. Condensation is quantified by the number of drops (green, left) and the total condensation area (red, right), both normalized to cell size, and compared between MinD-WT and FUS-IDR-MinD. (E) *In vitro* characterization of condensate-forming IDRs. Schematic of polySH3–IDR fusion constructs containing DDX4(1–235), LAF-1(1–228), FUS(2–214), and the droplet-deficient mutant FUS-Y27S(2–214) used for *in vitro* phase-separation assays (top). Representative fluorescence images of condensates formed by each construct in the polySH3–polyPRM reconstitution system are shown (bottom left). Scale bar: 10 μm. Quantification of droplet size distributions under identical conditions is shown on the bottom right: DDX4 and FUS formed large, coalesced condensates, whereas LAF-1 formed smaller condensates, and FUS-Y27S failed to undergo detectable phase separation. (F) Low-band-pass filtering of MinD-IDR coupling. Quantification of normalized condensate number (left) and droplet area (right) as a function of relative E/D expression level for different IDRs.

Using this approach, we first evaluated a D-FUS fusion RIPPLE system initially prototyped in our recruitment assays. Within the population, cells displayed a remarkable diversity of higher-order behaviors, ranging from wildtype-like MinDE waves and traveling condensed waves, to localized droplet arrays and highly condensed macrostructures (Figure 2B-C, Movie S3). Notably, even the most highly condensed, seemingly static structures remained actively driven, as we observed the spontaneous dissolution and reassembly of these structures coincident with characteristic spikes in local MinE levels.

Intriguingly, the transition between dynamic and condensed regimes appeared strongly correlated with the cellular MinE Activator expression levels, which tune wave speed. To systematically quantify this relationship, we profiled expression levels, oscillation frequencies, and condensation metrics (droplet number and total area) across thousands of individual cells and compared them to a control population expressing wildtype MinDE. As expected, the wildtype components yielded no detectable condensation at any expression level. In contrast, the FUS-MinD/E population exhibited robust condensation across a wide region of expression space.

Mapping this condensation as a direct function of the underlying reaction-diffusion waveform revealed a kinetic threshold: the system only permitted condensation at slow driving frequencies (<15 mHz; Fig. 2D). Importantly, these effects appeared to be specifically tied to the FUS IDR sequence’s ability to undergo phase separation, as a FUS-Y27S mutant known to disrupt droplet formation did not produce RIPPLE condensates at any expression level^58,59^. Thus, the D-FUS ATPase RIPPLE system appears to generate unique architectures in which protein condensation is gated by non-equilibrium wave dynamics.

To determine whether these RIPPLE behaviors reflect a general biophysical principle of coupling reaction–diffusion dynamics to condensation, we tested the ATPase-IDR RIPPLE platform using other well-characterized droplet-forming sequences, including DDX4 and LAF-1 (Fig. 2E-F; Fig. S2)^60,61^. As a reference point, we first benchmarked the intrinsic equilibrium droplet-forming propensity of these sequences using an engineered polySH3–polyPRM recruitment scaffold^39,41^. In this equilibrium setting, both FUS and DDX4 robustly formed large droplets with low saturation thresholds, while LAF-1 formed significantly smaller droplets, and the FUS-Y27S mutant failed to condense entirely (Fig. 2E; Fig. S2).

We then deployed these alternative sequences in our D-IDR RIPPLE framework and scored their behavior across thousands of unique cellular configurations. Like FUS, the D-DDX4 RIPPLE library generated dynamic, activity-driven condensation (Fig. S3). In contrast, the weaker LAF-1 sequence failed to condense under any wave conditions. Strikingly, the D-DDX4 RIPPLEs exhibited a distinct kinetic profile from D-FUS, with DDX4 condensates vanishing at significantly lower driving frequencies (5 mHz vs. 15 mHz). This divergence suggests differences in the underlying kinetic constraints on condensation of these IDRs are manifest in the non-equilibrium setting of RIPPLE. Nevertheless, both D-FUS and D-DDX4 exhibited the same qualitative dependence on wave dynamics, with condensation restricted to a low-frequency regime. Thus, coupling droplet-forming IDRs to the MinD ATPase appears to impose a dynamic low-pass filtering mechanism in which reaction–diffusion wave speed gates phase separation, and the intrinsic properties of an IDR determine the position of the condensation threshold.

### Frequency bandpass dynamics pattern droplet lattices in E-IDR Activator RIPPLEs

Targeting IDRs to the MinE-Activator at the rear of the wave inverts the spatial architecture of RIPPLE, fundamentally altering the relationship between reaction-diffusion effects and condensation. Instead of the continuous, condensed wavefronts seen in ATPase-IDR fusions, E-IDR targeting produces discrete, spherical droplets that desorb from the trailing edge of the wake and become responsive to wave dynamics. To define the rules governing this regime, we charted the phenotypic landscape of RIPPLE libraries sampling different E-IDR sequences across randomized expression levels in cells (Fig. 3, Movie S4).

**Figure 3:**
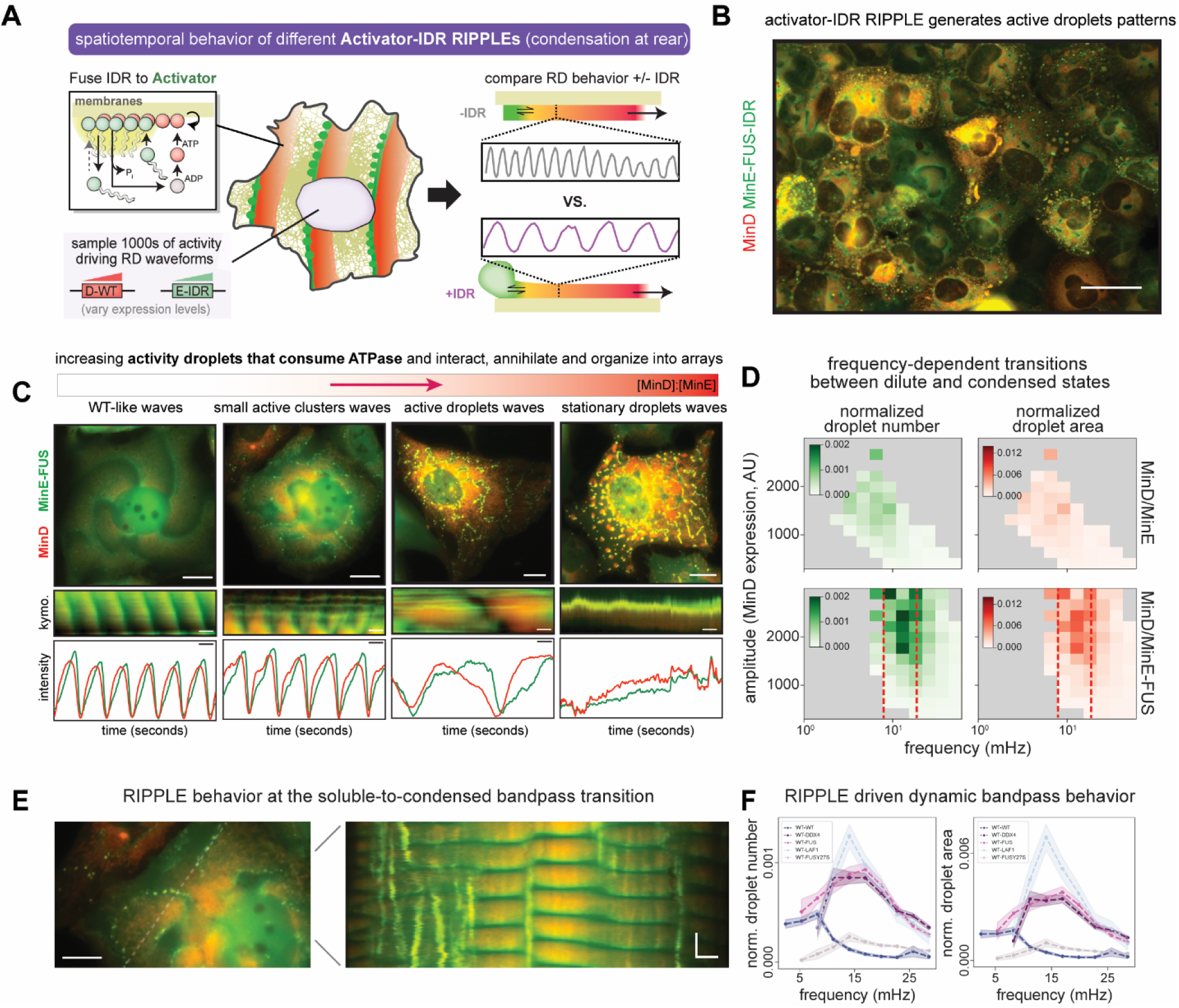
Activator-IDR coupling reveals band-pass filtering of condensate assembly by reaction–diffusion dynamics. (A) Schematic of constitutive MinE-IDR RIPPLE design. IDR sequences are directly fused to MinE and co-expressed with wild-type MinD. (B) Representative fluorescence images of MinE-FUS-IDR expressing cells across a population, displaying diverse spatial organizations. Scale bar: 50 μm. (C) Representative single-cell images illustrating the transition of condensation behavior as a function of relative MinD-to-MinE (D/E) expression ratio. Increasing MinD levels drive a shift from WT-like oscillatory patterns to active propagating droplets, and ultimately to persistent, spatially dominant condensation. Scale bar: 20 μm. (D) Two-dimensional binned histogram of population condensation score. Condensation is quantified by the number of drops (green, left) and the total condensation area (red, right), normalized by cell size, and compared between MinE-WT and MinE-FUS-IDR. (E) Representative high-magnification fluorescence image (left) and associated kymograph (right) of cells expressing MinE-FUS-IDR, capturing the transition from dynamic droplets to persistent condensates at the trailing edge of the wave patterns. Scale bar: 15 μm (fluorescence image). Kymograph scale bars: 5 μm (horizontal distance) and 30 s (vertical time). (F) Band-pass filtering of MinE-IDR coupling. Quantification of normalized condensate number (left) and droplet area (right) as a function of relative E/D expression level for different IDRs.

For the E-FUS RIPPLE system we had prototyped in our recruitment assays, direct fusion produced a dazzling array of cellular droplet patterns, networks, and wave phenomena that depended on both MinD and MinE expression level (Fig. 3B-C, Movie S4). Because MinD provides the membrane platform for MinE assembly and controls wave amplitude, droplets were seldom observed at low MinD expression levels, even when E-FUS expression was high and robust reaction-diffusion waves were seen. As MinD expression increased, however, spherical droplets of MinE began to appear, typically localizing to the spiral center or nodal structure of the reaction-diffusion field. When MinE-FUS and MinD expression were both high, individual droplets coalesced into highly ordered, evenly spaced labyrinthine macrostructures that decorated the cell interior. Intriguingly, these continuous networks behaved as physical boundaries to wave propagation, funneling and directing the flow of MinDE wave traffic throughout the cell interior.

Quantification of the condensation landscape revealed that droplet formation scores were significantly lower compared to the ATPase-IDR RIPPLE, consistent with MinE-IDRs favoring production of discrete droplets as opposed to larger condensed wavefronts. Mapping condensation metrics onto the underlying reaction-diffusion waveform and expression space revealed a distinct kinetic signature: while the ATPase-IDR platform operates as a low-pass filter, the Activator-IDR platform behaves as a narrow frequency bandpass filter (Fig. 2D; Fig. 3D). For E-FUS, robust condensation peaked at 17 mHz and dropped off when the underlying dynamics were faster or slower. This “Goldilocks zone” likely reflects a kinetic tension unique to rear-targeted RIPPLE architectures: high levels of E-FUS are required to drive local concentration past the condensation threshold, but these elevated activator levels drive the underlying MinDE reaction-diffusion engine to higher frequencies too fast for droplet formation.

To explore the generality of this bandpass behavior and its relationship to IDR sequence composition, we deployed the model DDX4, LAF-1, and non-condensing FUS-Y27S mutant IDRs in this E-IDR Activator RIPPLE context (Fig. 3F; Fig. S4 and S8; Movie S4). All E-IDR Activator sequences, with the exception of the FUS-Y27S mutant, generated and patterned droplets within the cell. However, each E-IDR sequence exhibited a unique, quantitatively distinct frequency bandpass profile. Surprisingly, the LAF-1 IDR, which exhibited weak, small droplet formation at equilibrium *in vitro* and failed to condense entirely when coupled to the ATPase front, was the strongest droplet-forming sequence in the E-IDR Activator RIPPLE context. This droplet formation activity, however, was restricted to a much sharper and narrower range of frequencies than either DDX4 or FUS, which supported weaker droplet formation across a wider range of waveforms.

This unexpected divergence highlights how out-of-equilibrium driving forces can override thermodynamic condensation hierarchies. Because LAF-1 condensates have been previously reported to possess higher molecular permeability and exchange kinetics than other model droplet-forming sequences, they may be better positioned to exploit the transient, high-flux environment of the MinDE wave tail^62,63^. However, this may come at the expense of robustness, restricting condensation to a narrow window of compatible driving waveforms. Collectively, these results demonstrate that E-IDR Activator RIPPLEs reveal kinetic and biophysical constraints that manifest distinctly from front-targeted D-IDR RIPPLEs—acting as a tunable bandpass filter that patterns discrete droplet lattices and wave barriers in the wake of the reaction-diffusion engine.

### Dual-IDR targeted ATPase/Activator RIPPLEs generate active condensate emulsions

The distinct phenomena elicited by coupling phase-separating sequences to either the front or the rear of a MinDE wave highlight how RIPPLE behavior is jointly determined by the intrinsic properties of the IDR and its point of spatial integration. This architecture raises the question of what emergent behaviors arise when IDRs are targeted simultaneously at both nodes of the reaction-diffusion circuit. Sustained propagation of wildtype MinDE waves depends on establishing distinct non-equilibrium spatial profiles of MinD and MinE across the total wavelength^25,52^. Conversely, different phase-separated protein condensates can exhibit varying degrees of mutual miscibility at equilibrium^64^. Dual-targeted D-IDR/E-IDR RIPPLEs, therefore, present an exotic non-equilibrium regime in which the reaction-diffusion engine’s intrinsic unmixing profile must interact with the biophysical rules governing condensate co-existence and phase separation (Fig. 4A).

**Figure 4:**
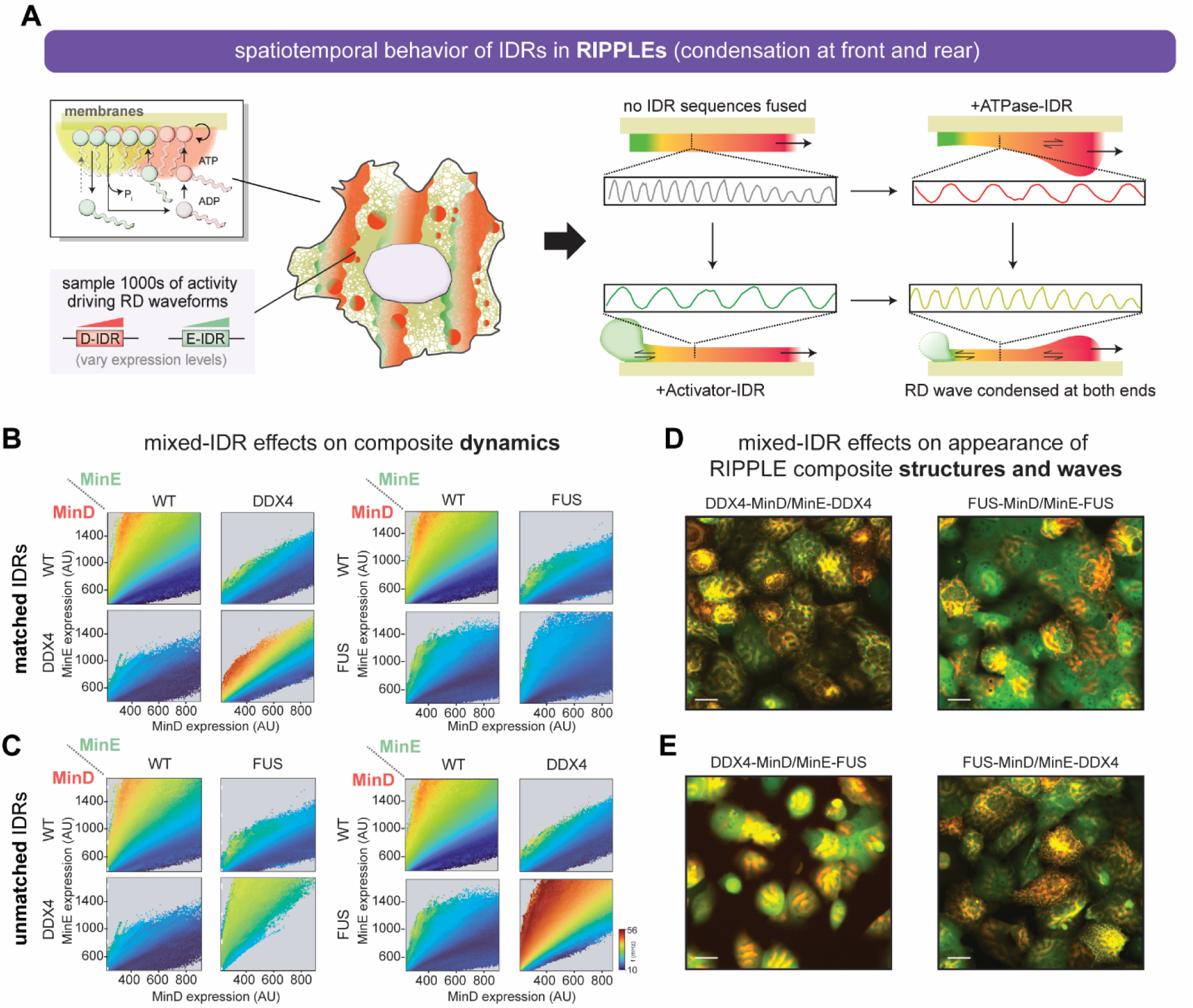
Dual-node IDR coupling rewires reaction–diffusion dynamics and emulsifies condensate organization. (A) Schematic of dual-node coupling in the MinDE circuit. IDRs are fused to MinD, MinE, or both nodes, enabling combinatorial control over phase separation and reaction–diffusion dynamics. Distinct coupling configurations modulate both cooperative and competing interactions between the two modules. (B) Effects of matched IDR coupling on MinDE signaling dynamics. Phase portraits of oscillation frequency across MinD and MinE expression space. Each 4×4 grid represents coupling configurations in which IDRs are fused to neither, either, or both nodes of the circuit. Left: DDX4; Right: FUS. (C) Effects of unmatched (orthogonal) IDR coupling on MinDE signaling dynamics. Same analysis as in (B), but with distinct IDRs fused to MinD and MinE. (D) Effects of matched IDR coupling on subcellular structural organization. Representative wide-field fluorescence images of matched IDR coupling at both nodes. Left: DDX4-IDR-MinD/MinE-DDX4-IDR; Right: FUS-IDR-MinD/MinE-FUS-FUS. Scale bars: 20 μm. (E) Effects of unmatched (orthogonal) IDR coupling on subcellular structural organization. Representative wide-field fluorescence images of unmatched IDR coupling. Left: DDX4-IDR-MinD/MinE-FUS-IDR; Right: FUS-IDR-MinD/MinE-IDR-DDX4. Scale bars: 20 μm.

To explore this interface, we characterized the steady-state behaviors of both matched (same IDR) and unmatched (different IDR) dual-RIPPLE libraries using the FUS and DDX4 sequences, which individually produced robust condensation phenotypes. Surprisingly, dual-IDR RIPPLEs exhibited qualitative and quantitative dynamics wholly distinct from their single-IDR counterparts (Fig. 4B-E; Movie S5). For both matched and unmatched dual-node configurations, the highly condensed macroscopic wavefronts characteristic of D-IDR ATPase RIPPLE and the discrete droplet arrays typical of E-IDR Activator RIPPLE were largely suppressed. Instead, these dual-node systems were dominated by extensive, persistent wave propagation across nearly all expression levels, suggesting that dual-targeting alters the physical constraints on condensation within the wave.

In the matched IDR combinations D-FUS/E-FUS and D-DDX4/E-DDX4, the waves propagated as a turbulent emulsion with a heterogeneous, fine-scale structure characterized by small, transient clusters and a high density of topological defects (Fig. 4D). Despite their overall morphological similarity, tracking the wave kinetics across matched expression levels revealed that these two configurations had opposing consequences on wave dynamics (Fig. 4B). While D-FUS/E-FUS waves propagated slower than wildtype MinDE, the D-DDX4/E-DDX4 combination produced faster waves. This was unexpected, given that the D-DDX4/E-WT RIPPLE and D-WT/E-DDX4 libraries both depressed wave speeds relative to wildtype controls. This suggests that homotypic IDR pairs allow the MinD and MinE machinery to mutually infiltrate the condensed phase, but variations in the underlying material properties, viscosity, or molecular diffusivity within these specific domains dictate how the resulting structure propagates through space.

Evaluating the unmatched heterotypic combinations (DDX4/FUS and FUS/DDX4) revealed an additional layer of complexity: the system exhibited orientation-specific behavior. For the D-DDX4/E-FUS RIPPLE configuration, wave propagation was virtually indistinguishable from wildtype, producing almost no detectable clusters or substructure (Fig. 4E). In contrast, the inverse D-FUS/E-DDX4 RIPPLE configuration displayed extensive cluster formation, transient micro-droplets, and a highly granular morphology during propagation (Fig. 4C). Thus, while dual-node targeting generally restores wave activity by allowing components spatial access to one another, the incomplete miscibility reported for DDX4 and FUS droplets at equilibrium manifests in a non-commutative fashion depending on where along the non-equilibrium waveform each sequence is positioned^65,66^.

Although these spatial asymmetries initially appear paradoxical, they are consistent with the distinct kinetic filtering thresholds observed in our single-IDR RIPPLE assays. Condensation of the more cohesive, kinetically stable FUS IDR at the ATPase front favors the initial formation of larger structural clusters^67,68^. However, because E-DDX4 can partition into these assemblies, it can stimulate ATP hydrolysis and wave propagation, breaking down condensates before large-scale macrostructures can stabilize. Conversely, when the more kinetically sensitive DDX4 sequence occupies the leading wavefront, the trailing MinE-FUS machinery infiltrates and dismantles developing condensates at an even earlier stage, preventing them from maturing into resolvable clusters^65,69^. Consequently, the macroscale concentration gradients maintained by reaction-diffusion impose asymmetric rules on the tug-of-war between co-existing phase-separated wave regions.

Together, these observations indicate that dual-node coupling within a reaction-diffusion circuit can drive an emulsified condensate state in which macroscopic phase separation is suppressed and replaced by dispersed, dynamically regulated micro-assemblies. Importantly, the exact spatial properties and dynamical consequences of this emulsification depend on the intrinsic material traits of the selected IDR sequences. RIPPLE thus demonstrates how reaction-diffusion transport can both constrain and be reshaped by phase separation, suggesting a combinatorial strategy for engineering complex intracellular organization by tuning distinct interaction modules across multiple positions within a shared dynamical circuit.

### Diverse RIPPLE IDR combinations encode a spectrum of active subcellular architectures

By coupling model droplet-forming IDR sequences like FUS and DDX4 to different nodes of a self-organizing reaction-diffusion (RD) engine, we have shown how RIPPLE generates emergent subcellular architectures, patterns, and emulsions. However, IDR sequences are widespread across evolution and have diversified extensively, imparting them with distinct biophysical and material properties and unique miscibility profiles. This raises the intriguing question: how does the same reaction-diffusion engine organize and react to diverse IDR inputs and combinations?

Using RIPPLE, we explored the phenotypic landscape spanned by a combinatorial library of diverse droplet-forming sequences: model IDRs (FUS and DDX4), eukaryotic RGG-rich domains (LAF-1 and GAR-1)^70^, and the highly diverged bacterial sequence PopZ^71–73^. Each IDR was tested as an isolated D-IDR ATPase or E-IDR Activator RIPPLE, as well as all pairwise D-IDR/E-IDR RIPPLE combinations (Fig. 5A; Fig. S5; Movies S6-8). This enabled us to probe how co-condensation and miscibility effects compete with, antagonize, or synergize with the underlying driving reaction-diffusion waveform. Across this combinatorial space, we observed two major behavioral regimes: one favoring persistent, ordered macrostructures (Fig. 5B) and another dominated by dynamic emulsification (Fig. 5C).

**Figure 5:**
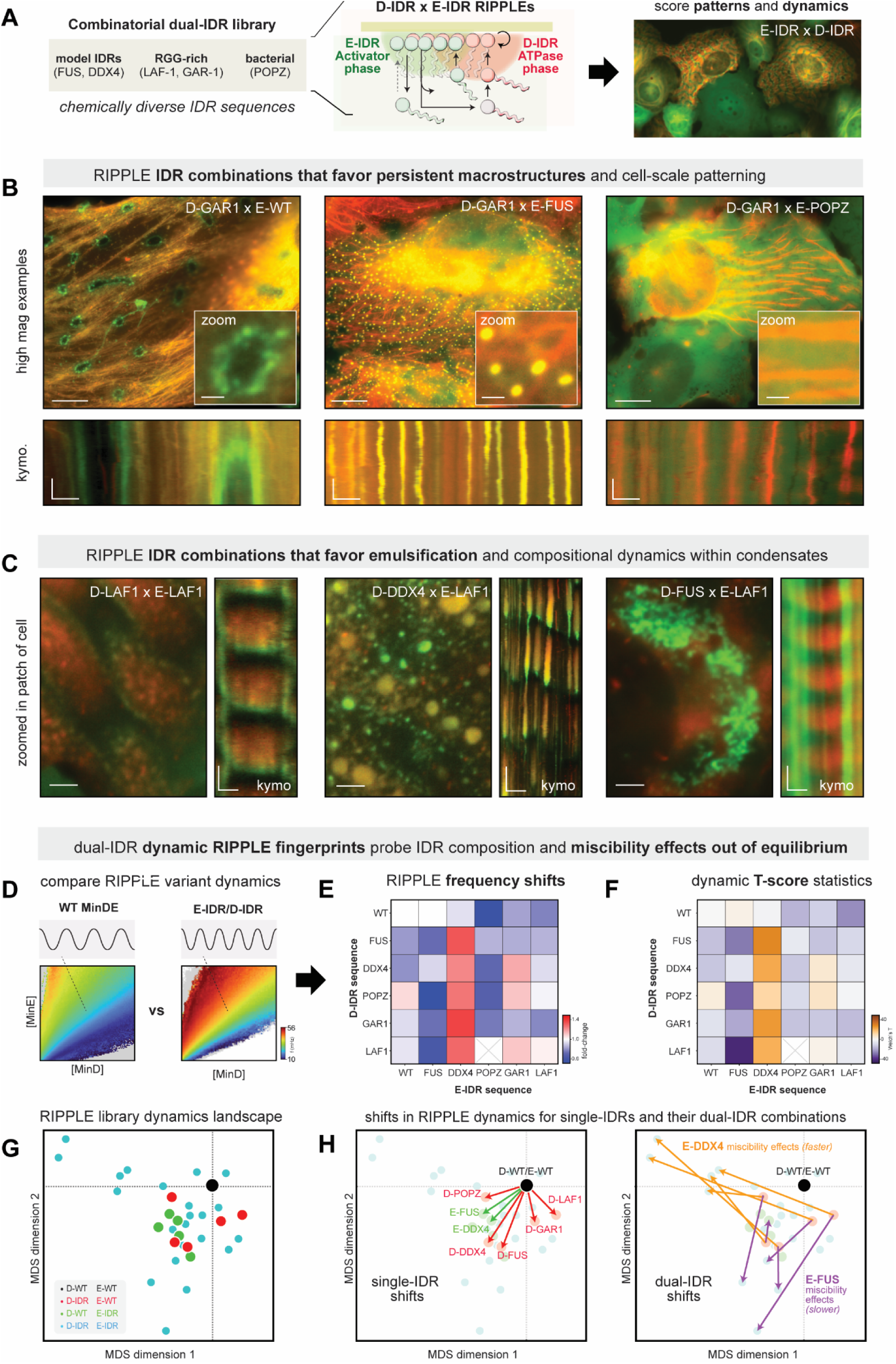
Orthogonal IDR coupling generates a diverse landscape of multiphase architectures and reaction–diffusion behaviors. (A) Schematic of the reaction–diffusion (RD)-condensation design space. Diverse IDR sequences are coupled to the MinDE circuit at distinct network nodes and sampled across MinD/MinE expression space, generating a combinatorial landscape of interaction strengths and dynamical regimes. (B) Representative fluorescence images showing persistent macrostructures with distinct cellular patterning generated by different RIPPLE IDR combinations. From left to right: GAR1-MinD/MinE, GAR1-MinD/MinE-FUS, and GAR1-MinD/MinE-POPZ. High-magnification views of the indicated regions are shown above. Scale bars: 15 μm (whole-cell images) and 2 μm (zoomed images). Corresponding kymographs are shown below. Kymograph scale bars: 3 μm (horizontal distance) and 3 s (vertical time). (C) Representative fluorescence images showing dynamic emulsification and compositional remodeling generated by different RIPPLE IDR combinations. From left to right: LAF1-MinD/MinE-LAF1, DDX4-MinD/MinE-LAF1, and FUS-MinD/MinE-LAF1. For each combination, a zoomed view is shown on the left (scale bar, 5 μm), with the corresponding kymograph on the right. Kymograph scale bars: 3 μm (horizontal distance). Vertical time scale bars are 2 s for LAF1-MinD/MinE-LAF1 and FUS-MinD/MinE-LAF1, and 6 s for DDX4-MinD/MinE-LAF1. (D-H) Dual-IDR RIPPLE fingerprints reveal the effects of IDR composition and miscibility on RD dynamics. (D) Schematic illustrating pairwise comparisons between dual-IDR RIPPLE variants and the baseline wild-type MinDE circuit. (E) Heatmap of oscillation frequency shifts relative to the baseline. (F) Heatmap of corresponding T-score states relative to the baseline. (G,H) Multidimensional scaling (MDS) analysis of oscillation dynamics across RIPPLE variants. (G) Dynamical landscape of all tested IDR combinations visualized by MDS. (H) MDS trajectories showing dynamical shifts associated with single-IDR fusions (right) and corresponding dual-IDR combinations (left). The distinct effects of MinE-DDX4 (orange) and MinE-FUS (purple) are highlighted.

In the first regime, stable subcellular architectures formed that arrested or attenuated reaction-diffusion dynamics (Fig. 5B; Movie S6). For example, D-GAR1 condensed so strongly along the endomembrane that it became difficult for the wild-type MinE activator to penetrate this platform^41,74^. This caused the MinE activator to treadmill, organizing into higher-order dissipative rings and stripes patterned at a consistent length-scale, and which excluded MinD from their interior. Pairing D-GAR1 with different E-IDR activators fundamentally reorganized this macroscopic patterning: E-POPZ fragmented the continuous D-GAR1 platform, remodeling it into ordered arrays of spear-like structures aligned with cell geometry; while E-FUS condensed directly on the dense D-GAR1 platform, collapsing the ring structures into more discretely patterned droplets. Importantly, these subcellular architectures were actively maintained by the underlying reaction-diffusion engine, as we occasionally observed the spontaneous appearance or disappearance of sub-structures within or along the overall cell-scale architecture.

Other RIPPLE-IDR combinations favored dissipative emulsification, creating stunning condensation behaviors with rich compositional dynamics (Fig. 5C; Movie S7). For example, while E-LAF1 Activator RIPPLE on its own resulted in droplet lattices, dual-IDR pairings led to a range of emulsion behaviors. The self-paired D-LAF1/E-LAF1 RIPPLE drove robust wave propagation across all expression levels, maintaining a highly fluid, fine-droplet emulsion visible only at high magnification. In contrast, D-DDX4/E-LAF1 RIPPLE produced remarkable condensates that oscillated in size and composition, cycling back and forth between large D-DDX4 droplets and small E-LAF1 droplets. Interestingly, D-FUS/E-LAF1 RIPPLE caused E-LAF1 aggregates to accumulate in the cell, even though faint low-amplitude reaction-diffusion waves continued to propagate around them. Because FUS and LAF1 are reported to have low co-miscibility, we hypothesize that dilute E-LAF1 molecules can condense during assembly of the D-FUS wavefront, but struggle to enter subsequent wave cycles once phase separated^64,65,75^. These results highlight the complexity of condensation and miscibility when sequences are strongly driven out of equilibrium, and the utility of the RIPPLE platform to probe and reveal these effects.

To gain quantitative insights into the underlying reaction-diffusion engine driving these phenomena, we scored each IDR combination’s effect on RIPPLE dynamics (Fig. 5D-F; Fig. S6). For single-targeted D-IDR or E-IDR RIPPLEs, reaction-diffusion dynamics were generally attenuated. Dual-targeted D-IDR/E-IDR RIPPLEs were also generally slower than wildtype MinDE, with specific exceptions associated with particular IDR sequences. For example, while single-targeted E-DDX4 RIPPLEs were slow on their own, E-DDX4 drove faster reaction-diffusion dynamics when paired with any other IDR sequence we tested, including sequences that dramatically slowed wave dynamics in all other IDR pairings (e.g., D-FUS, D-GAR1). For E-GAR1, dual-IDR RIPPLEs accelerated dynamics in some contexts, but slowed in others. In contrast, E-FUS further attenuated the wave dynamics of all other D-IDR sequences except for D-DDX4.

To visualize these combinatorial effects on dynamics across the entire library, we used the frequency shift between any two RIPPLE systems as a distance metric and used multidimensional scaling (MDS) to organize the library into a compact two-dimensional space, where points close in space indicate similar dynamic behaviors and RIPPLE miscibility profiles^76^ (Fig. 5G-H; Fig. S7). Here, we can see that pairing E-DDX4 with D-IDRs, but not D-WT, pulls the RIPPLE system into a specific region of the MDS space associated with faster dynamics, while E-FUS pairings pull the system in the opposite direction towards slower dynamics. Thus, our combinatorial RIPPLE screen not only reveals a rich diversity of subcellular architectures but also provides a dynamic fingerprint of how an IDR sequence’s biophysical properties manifest when challenged to operate in an out-of-equilibrium, multi-phase context.

### RIPPLE architectures provide a platform for the recruitment of other protein components

In natural systems, structured subcellular architectures provide a platform for the assembly of other macromolecules^40,42^. To test whether RIPPLE architectures were compatible with protein recruitment, we generated FRB-tagged RIPPLEs that operated in either a dynamic, condensed wavefront regime (D-FUS ATPase RIPPLE) or patterned droplet lattice regime (E-FUS Activator RIPPLE), and then tested their ability to co-condense an FKBP-BFP payload upon recruitment with rapamycin (Fig. 6A; Movie S9)^57^.

**Figure 6:**
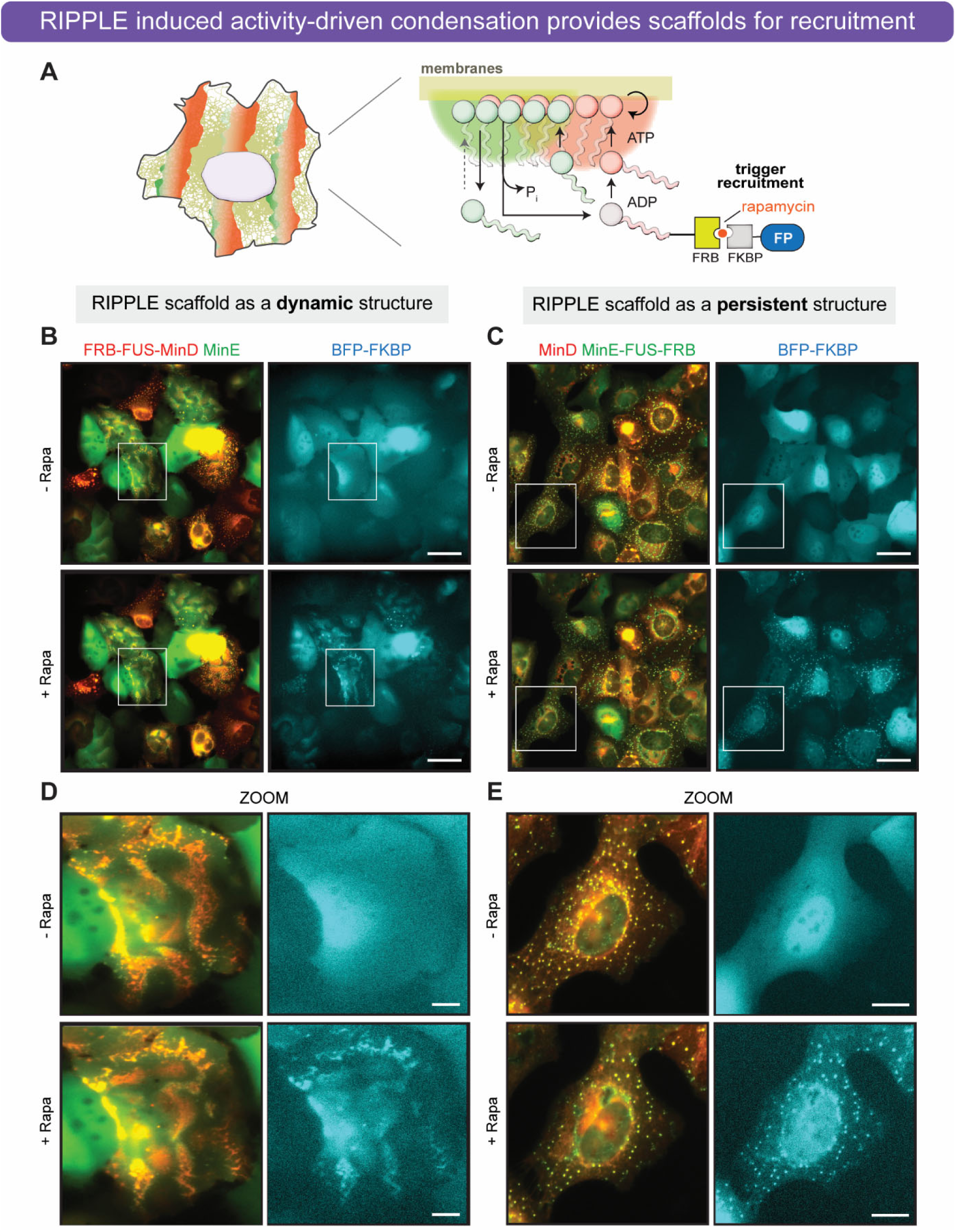
RIPPLE architectures can scaffold the assembly of other macromolecular clients. (A) Schematic of inducible client recruitment in the RIPPLE system. FRB-tagged MinD/MinE constructs fused to FUS-IDR are co-expressed with a soluble BFP-FKBP reporter. Addition of rapamycin induces FKBP–FRB heterodimerization, enabling recruitment of the reporter into condensate structures formed within the reaction–diffusion circuit. (B) Representative widefield images of FRB-FUS-IDR-MinD before and after induction. Prior to rapamycin addition, BFP-FKBP is diffusely distributed and uncoupled from MinD dynamics. Upon induction, the reporter is recruited to FUS-IDR-enriched regions, resulting in localization to condensate-associated structures and partial synchronization with the underlying wave patterns. Scale bars: 50 μm. (C) Representative widefield images of MinE-FUS-IDR-FRB before and after induction. Similar inducible recruitment is observed when the FRB module is positioned at the MinE node, demonstrating that activity-driven client localization can be achieved across distinct circuit integration points. Scale bars: 50 μm. (D) Zoomed view of the recruitment behavior shown in (B), highlighting localization of BFP-FKBP within dynamic RIPPLE condensate structures associated with propagating wave patterns. Scale bars: 10 μm. (E) Zoomed view of the recruitment behavior shown in (C), highlighting recruitment of BFP-FKBP into persistent RIPPLE condensate structures. Scale bars: 20 μm.

In the absence of inducer, the FKBP-BFP signal was diffuse throughout the cytoplasm for both the D-FUS ATPase and E-FUS Activator RIPPLEs (Fig. 6B-E). The addition of rapamycin led to rapid recruitment and assembly of the BFP payload in both RIPPLE systems. For the D-FUS ATPase RIPPLE, the BFP payload became synchronized with RIPPLE activity, exhibiting cycles of enrichment and dispersal coordinated with assembly and disassembly of the condensed wavefront; and for E-FUS Activator RIPPLEs, the BFP payload became concentrated directly into the droplet lattice. Thus, activity-driven condensates and architectures generated by RIPPLE can serve as a scaffold for recruiting other client proteins, mirroring the control logic for client recruitment into passively formed condensate scaffolds ^40^. RIPPLE thus provides a foundation for engineering programmable intracellular organization dynamically regulated by an underlying reaction-diffusion circuit.

## Discussion

By coupling a programmable reaction-diffusion engine to droplet-forming IDR sequences, RIPPLE provides a synthetic platform for understanding and engineering active subcellular architectures and protein condensation far from equilibrium (Fig. 7). The reaction-diffusion arm is provided by MinDE protein waves that propagate along membranes with tunable amplitude and frequency, driven by assembly of a MinD-ATPase front disassembled at its rear by the MinE-Activator. Connecting droplet-forming IDR sequences to different components of this wave circuit allows these rapidly changing local concentrations to drive condensate assembly and disassembly in different ways. Feedback between condensation and wave dynamics generated a spectrum of surprising subcellular architectures, including condensation waves, droplet networks, dynamic emulsions, and phase-separated macrostructures. The output structure was a product of both IDR sequence composition and miscibility, as well as the underlying RD waveform dynamics, with frequency-dependent transitions regulating the transition between dilute and condensed states. The RIPPLE platform thus provides a non-equilibrium molecular multitool for understanding and engineering protein condensation out of equilibrium, and for exploring the spectrum of subcellular architectures that can arise when reaction-diffusion signaling and phase separation intersect.

**Figure 7.**
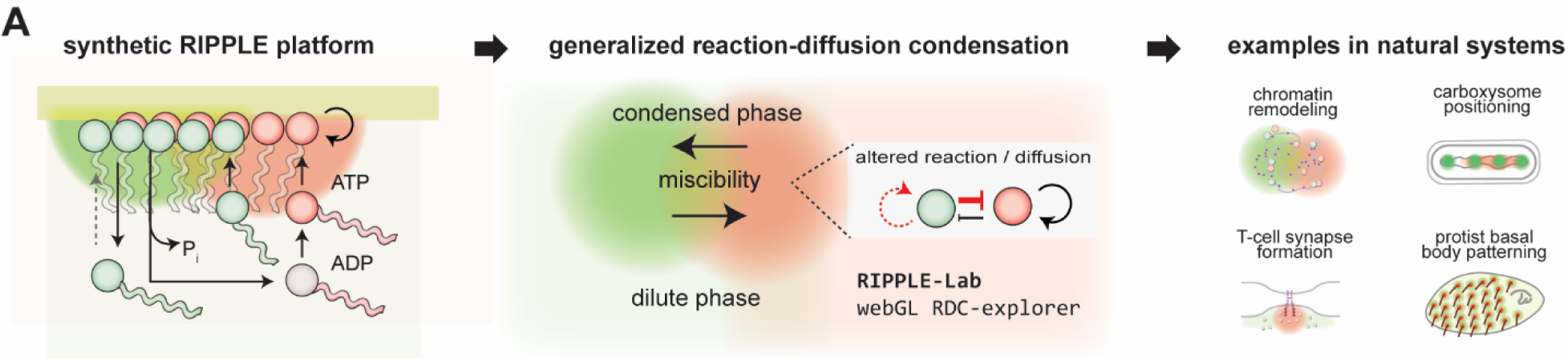
Reaction–diffusion-condensation as a framework for regulatable active subcellular architectures. Conceptual framework for reaction–diffusion-condensation (RDC) coupling. RIPPLE provides a minimal two-protein system in which reaction–diffusion dynamics and protein condensation interact to generate diverse subcellular architectures. We propose that similar RDC principles may operate across natural cellular systems that couple signaling and structure.

### RIPPLE probes protein condensation and IDR sequence behavior far from equilibrium

Our core understanding of protein condensation is largely derived from equilibrium studies: droplets form when protein concentration exceeds the saturation concentration *c_sat_*^77^. However, cellular condensates interface with active signaling networks that use energy to continuously remodel molecular composition and organization. This activity-driven regulation is thought to be critical for a range of cell-biological functions, including chromatin remodeling, gene regulation, and commitment to T-cell synapse formation during target recognition^45,47^. In these non-equilibrium contexts, dynamic changes in local protein concentration near *c_sat_* interact with condensation timescales, placing the system’s behavior in a kinetically controlled regime. Previous efforts to explore this regime in cells using optogenetic triggers or *in vitro* using motor protein assemblies have revealed novel kinetic effects on condensation^44,78–80^. However, systematic investigations are constrained by throughput and limitations on tunable dynamic control. Incorporating multiple interacting phases in these systems is even more challenging.

In contrast, RIPPLE provides a fully programmable reaction-diffusion engine that can subject a wide spectrum of dynamics to IDR sequences both in isolation or in mixtures across thousands of cells in parallel. This scalable platform allowed us to systematically explore the relationship between the RIPPLE waveform and condensation for any IDR sequence, revealing frequency-dependent effects that varied across IDR sequences. Dual-targeting IDR sequences allowed exploration of how miscibility can manifest asymmetrically in an out-of-equilibrium context, with strongly miscible sequences driving emulsification and less miscible sequences favoring active exclusion into phase-separated patterns. By testing multiple IDRs across many pairwise combinations, RIPPLE generates dynamic multi-feature fingerprints for IDR sequences that complement equilibrium metrics like *c_sat_*. Thus, RIPPLE enables exploration of IDR sequence effects that are likely to be relevant in the out-of-equilibrium, multi-phase context of living cells.

### Reaction-diffusion-condensation as a framework for structure-signaling composites

The RIPPLE platform shows how reaction-diffusion and condensation can cooperate in a minimal two-protein system, but are the effects we see in this synthetic system representative of a broader paradigm relevant to other naturally evolved systems? Although the detailed kinetic schemes at play in MinDE-IDR RIPPLE systems are complex^25,37,49,53–56,81–83^, the key features modulating RIPPLE behavior arise from two core mechanisms. First, the underlying reaction-diffusion engine cyclically drives condensation by increasing and decreasing local IDR concentrations. Within these condensed ATPase or Activator phases, any parameter of the reaction-diffusion mechanism could be modulated, including binding, catalysis, and diffusion. Second, when different IDR sequences drive ATPase and Activator into distinct chemical phases, the concentration gradient imposed by the reaction-diffusion engine and the preferred compositional profiles of the associated phases can begin to interact in non-trivial ways.

These effects—modulation of reactivity and diffusivity within a condensate and miscibility effects—would seem generalizable to many natural circuits that couple structure and signaling together in cells. To gain intuition, we extended a widely studied Gray-Scott Reaction-Diffusion system^84^ to allow inclusion of these condensation effects and implemented a real-time RIPPLE-lab RDC explorer webGL simulation that enables rapid exploration of parameter space (https://coylelab-uw-madison.github.io/RIPPLE-RDC-explorer/). Starting from reaction-diffusion parameters that generate propagating waves, condensation-dependent parameter modulation, as well as miscibility effects, can expand the phenotypic landscape accessible to the simulation, with direct parallels to many observations made experimentally with RIPPLE. The resulting phenotypes included droplet formation that could be emulsified by increasing co-miscibility, and oscillating droplet networks arising from modulation of droplet reactivity and diffusivity.

Our toy model points to reaction-diffusion-condensation (RDC) as a more general mechanism for generating composite structure-signaling systems with a rich and tunable phenotypic landscape. We hypothesize that RDC mechanisms are likely at play across a range of natural cellular settings. RDC mechanisms can act locally, allowing dynamics and activity to regulate the structure of individual condensates or assemblies. We can see such logic in the formation of the T-cell synapse, where kinase/phosphatase signaling controls the recruitment and local concentration of proteins that undergo a phase transition to lock in target selection via phosphatase exclusion; as well as in gene-regulatory hubs, where cycles of chromatin remodeling and transcriptional activity inject energy into transcriptional condensates^45,74^. Importantly, because the underlying reaction-diffusion signaling dynamics are required to maintain these active structures, cells can rapidly regulate their formation, composition, and lifetime to build responsive living assemblies.

Interestingly, RDC mechanisms can also cooperate over longer length scales, allowing local structural regulation to propagate into whole-cell patterns, macrostructures, and subcellular architectures. Indeed, such higher-order self-organizing mechanisms are thought to be at play in the Rho-Actin waves seen in marine embryos and certain immune cell types, which connect endogenous reaction-diffusion waves of GTPase signaling to the cytoskeleton; and in the reaction-diffusion-driven positioning of plasmids and carboxysome condensate arrays in cyanobacteria^23,85–89^. More provocatively, the diverse and tunable spectrum of patterns and subcellular architectures we generated synthetically bears a striking resemblance to the plethora of cortical patterns of basal bodies, motile cilia, and feeding structures seen across protists whose mechanistic origins have bewildered cell biologists for over a hundred years^12,18,90–93^. While such systems are almost certainly refined and made more complex over the course of evolution, RIPPLE suggests that the starting point for these structures may be realized more simply: a minimal two-protein circuit integrating structure and signaling is sufficient to unlock a tunable and regulatable landscape of seemingly endless subcellular architectures.

## Acknowledgements.

We thank members of the Coyle Lab, A. Weeks and W. Lim for advice, helpful discussions, and critical reading of the manuscript.

## Author contributions

Conceptualization EC, ZX and SMC; methodology EC, ZX, EWZW and SMC; Investigation EC, ZX, EWZW and SMC; writing – original draft, EC, ZX and SMC; writing – review & editing EC, ZX, EWZW and SMC; funding acquisition, SMC; resources SMC.

