## Supplemental Materials and Methods for "Reaction-diffusion condensation generates a regulatable landscape for self-organizing subcellular structures"

### Supplementary materials for:
### Reaction-diffusion condensation generates a regulatable landscape for self-organizing sub-cellular structures

Eden Chang[1,2,*,‡], Zhejing Xu[1,3,*], Elliott W. Z. Weix[1], Scott M. Coyle[1,#]

**Affiliations:**
[1] Department of Biochemistry
[2] Biophysics Graduate Program
[3] Integrated Program in Biochemistry Graduate Program

[*] These authors contributed equally to this work

[‡] Present address: UT Southwestern

University of Wisconsin-Madison, Madison, Wisconsin 53706, USA.

**This supplement contains:**

Materials and Methods
Supplementary Figures S1-8
Supplementary Video Legends S1-9

**Material and Methods.**

**Plasmid construction**

DNA fragments encoding mCh-MinD and MinE-EGFP were amplified by PCR using Addgene plasmids (#213112 and #213113) and subsequently subcloned into SFFV or Lenti expression vectors. To create IDRs-mCh-MinD and MinE-EGFP-IDRs, we used the following IDRs: Homo sapiens DDX4 (residues 1-236), FUS (residues 1-214), the FUSY27S mutant (residues 1-214), and LAF-1 RGG (residues 1-200). Additionally, we included the *C. elegans* GAR1 IDR (residues 1-67) and *E. coli* POPZ (residues 24-102) fragments, which were obtained as gBlocks from Integrated DNA Technologies. These IDR fragments were fused to the N-terminus of SFFV-mCh-MinD or the C-terminus of SFFV-MinE-EGFP. For $SH3_3$-IDRs and EGFP-$PRM_4$ constructs, IDRs were amplified by PCR using the templates described above and integrated into pET-28a(+) plasmids. The $SH3_3$ and $PRM_4$ fragment, obtained as gBlocks from Integrated DNA Technologies, was inserted into a pBH4 vector, which contains a monoGFP sequence and a His-tag at the N-terminal region.

**Lentiviral transduction of mammalian cell lines**

Pantropic VSV-G pseudotyped lentivirus was produced by transfecting 293T cells (ATCC CRL-3216) with SFFV or Lenti expression vectors and the viral packaging plasmids psPAX2 and pMD2.G using Fugene HD (Promega #E2312). Viral production was carried out in 6-well tissue culture-treated plates (Corning 3335). After 72 hours, the viral supernatant was harvested, filtered through a 0.45 μm PES syringe filter, and added to mammalian cell lines along with 2 μg/ml Polybrene transfection reagent (Sigma TR-1003-G). Viral medium was replaced with normal growth medium 24 hours after infection.

**Live cell imaging**

Cells were plated on glass-bottom plates (Cellvis P06-1.5H-N) to a confluency of 60–80% in FluoroBrite DMEM (ThermoFisher A1896701). After 1 day, cells were imaged on a Nikon Ti-Eclipse in a Tokai Hit stage-top incubator: 37 °C and 5% $CO_2$ conditions. Multi-channel fluorescence images were collected at 1 frame per second with a 50% intensity and 500 ms exposure time for each channel.

**Cell segmentation analysis**

Individual cells were segmented from time-averaged fluorescence images generated by combining channels. Segmentation was performed using the Cellpose algorithm (Pachitariu et al., 2025; Stringer et al., 2021), which assigns a unique label to each cell and generates corresponding cell contours. For each segmented cell, fluorescence intensities were quantified by averaging pixel values within the cell contour across both space and time. Specifically, the mean fluorescence intensity of IDRs-mCherry-MinD and MinE-EGFP-IDRs was calculated by averaging all pixels within each cell contour over the full time series.

**Fast-Fourier transform-based frequency-domain image analysis**

Time-lapse fluorescence images were analyzed in the frequency domain using the Fast Fourier Transform (FFT). All analyses were performed using custom Python scripts. The processing pipeline followed previously published methods (Rajasekaran et al., 2024) and is summarized here. For each pixel, the fluorescence intensity time series was first zero-centered by subtracting a fitted low-order polynomial to remove slow baseline decay. The zero-centered pixel-wise signals were then transformed from the time domain to the frequency domain using FFT. For each pixel, frequency components corresponding to bins 0–40 were retained (rather than the full $N/2$ range, where $N$ is the number of time points) to construct a truncated frequency-domain representation.

The magnitude (absolute value) of the complex FFT output was computed to obtain the power spectrum for each pixel.

For single-cell analysis, pixel-wise power spectra were normalized across retained frequency bins to minimize intensity-dependent effects. The frequency corresponding to the maximum normalized power was assigned as the dominant oscillation frequency for each pixel when the normalized power exceeded 0.30; pixels below this threshold were excluded. Dominant frequency and normalized power values were then aggregated across all retained pixels within each segmented cell contour to calculate cell-level mean oscillation frequency and mean normalized power for each fluorescence channel. To determine the contribution of wave dynamics to condensate formation across different RIPPLE designs, MinD and MinE expression levels were projected onto a reference oscillation axis derived from the baseline mCherry-MinD/MinE-EGFP condition. Previous work demonstrated that oscillation frequency is primarily governed by the MinE-to-MinD stoichiometric ratio, with an approximately linear dependence across the low-frequency regime examined here (Rajasekaran et al., 2024). Accordingly, the relationship between oscillation frequency and the MinE-to-MinD expression ratio was determined from the baseline phase portrait by linear regression. The resulting fit was then used to convert the measured MinE-to-MinD ratio into a predicted theoretical frequency, representing the oscillation frequency expected solely from MinD/E reaction-diffusion signaling dynamics.

For population-level analysis, additional quality-control filters were applied to ensure robust mapping between protein expression and oscillatory dynamics. Pixel-wise FFT spectra were retained only when both MinD and MinE signals exhibited a dominant peak with a signal-to-noise Z-score greater than 5 and contained a single detectable frequency peak. Retained pixels from each experimental condition were binned according to MinD and MinE expression levels to generate two-dimensional phase portraits, where the mean oscillation frequency was computed for each expression bin to characterize population-level wave dynamics across expression space.

**Pairwise wave dynamic comparison**

To compare wave dynamics across conditions, pairwise differences between phase portraits were quantified using bin-wise Welch's t-statistics. For each overlapping expression bin, a Welch's t-statistic was calculated from the corresponding frequency distributions, providing a measure of local differences in oscillatory behavior between conditions. A global difference metric was then computed by summing the squared t-statistics across all overlapping bins. Pairwise difference metrics were calculated for all phase portraits to generate a distance matrix describing the similarity of wave dynamics across conditions. The resulting distance matrix was projected into a two-dimensional space using non-metric multidimensional scaling (MDS) for visualization. MDS embeddings were generated using a fixed random seed of 42 to ensure reproducibility. Similar comparisons were obtained using alternative distance metrics, including Cohen's d and ΔFreq.

$$d_{cohen} = \frac{\overline{X_1} - \overline{X_2}}{s_{pooled}} \qquad T_{Welch} = \frac{\overline{X_1} - \overline{X_2}}{\sqrt{\frac{s_1^2}{n_1} + \frac{s_2^2}{n_2}}} \qquad \Delta freq = \overline{f_1} - \overline{f_2}$$

**Protein condensation score analysis**

To quantify condensate formation across IDRs-mCherry-MinD and MinE-EGFP-IDRs conditions, time-averaged fluorescence images were analyzed using a custom Python pipeline to detect localized, high-contrast structural foci within individual cells. To isolate condensates from diffuse cytoplasmic signals while minimizing artifacts at steep intensity gradients (such as the nuclear envelope), images from each channel were first background-subtracted using a local median filter (kernel footprint = 5 × 5 pixels). To eliminate intensity-dependent effects caused by variations in overall expression level, the resulting background-subtracted images were pixel-by-pixel normalized to their corresponding raw, unblurred time-averaged images, yielding a local contrast map representing the relative percentage drop in intensity. These normalized maps were empirically thresholded at a relative cutoff of 0.10 to generate binary masks of true condensate regions independent of absolute pixel values. At the single-cell level, condensates were quantified within segmented cell contours by measuring (i) the number of discrete condensate regions and (ii) the total condensate area. Final condensation scores were calculated by normalizing these metrics to the total cell area, providing a robust, per-cell measure of condensation propensity.

**Protein expression and purification**

All $SH3_3$-IDRs and EGFP-$PRM_4$ constructs were expressed in BL21(DE3). Bacteria were grown in LB at 37 °C and induced at A600 0.6-1.0 and expressed at 16 °C for 18 hours after induction with 0.5 mM IPTG. The bacterial culture was harvested by centrifugation and lysed according to the following procedure. The cell pellets were resuspended in lysis buffer (50 mM KH2PO4, 50 mM Na2HPO4, 150 mM NaCl, and 2 mM β-mercaptoethanol), disrupted by French press, and centrifuged at 15,000 g at 4°C for 30 min. The supernatant was incubated with Ni resin (Sigma) for 30 min, followed by a prewash with 25 mM imidazole. Proteins were eluted with the buffer containing 250 mM imidazole. The $His_6$ tag of $SH3_3$-IDRs was cleaved by TEV protease. The proteins were further purified using a MonoQ 5/50 anion exchange column. The samples were then subjected to size-exclusion chromatography (SEC) on a Superdex 200 16/60 column and analyzed by SDS-PAGE. For EGFP-PRM4 protein, the His6 tag was removed by TEV protease. The protein was then purified using a HiTrap S cation exchange column, followed by size-exclusion chromatography (SEC) on a Superdex 200 16/60 column. The peak protein fraction was collected, concentrated, and stored in a buffer of 150 mM KCl, 10 mM imidazole, 1 mM EGTA, 1 mM MgCl2, and 1mM DTT (pH 7.0). All proteins were flash-frozen and stored at -80 °C.

***In vitro* droplet formation assay**

The droplets were formed by mixing different $SH3_3$-IDRs with EGFP-$PRM_4$ at varying module concentrations in a buffer consisting of 150 mM KCl, 10 mM imidazole, 1 mM EGTA, 1 mM $MgCl_2$, and 1 mM DTT (pH 7.0). They were then visualized using Nikon AXR confocal microscopy. Images were measured and quantified using Fiji.

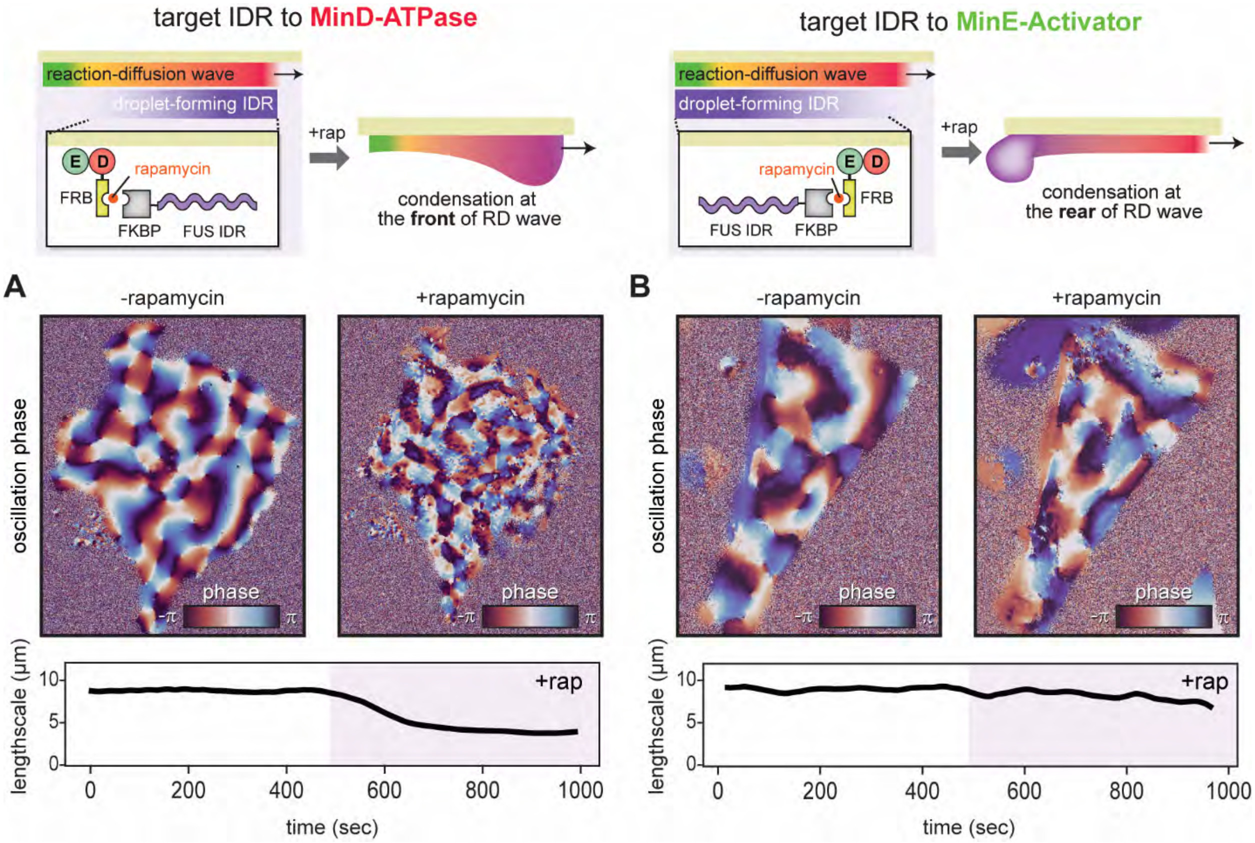


**Figure S1. Rapamycin-inducible RIPPLE modulation of reaction–diffusion wave organization.**

(A) Recruitment of FUS-IDRs to the ATPase MinD. Top: spatial phase maps derived from wave dynamics before (left) and after (right) rapamycin induction. Bottom: Quantification of characteristic spatial length scales extracted from the phase maps, showing remodeling of wave organization following induction.

(B) Recruitment of FUS-IDRs to the activator MinE. Top: spatial phase maps derived from wave dynamics before (left) and after (right) rapamycin induction. Bottom: Quantification of characteristic spatial length scales extracted from the phase maps, showing comparatively minor changes in wave organization following induction.

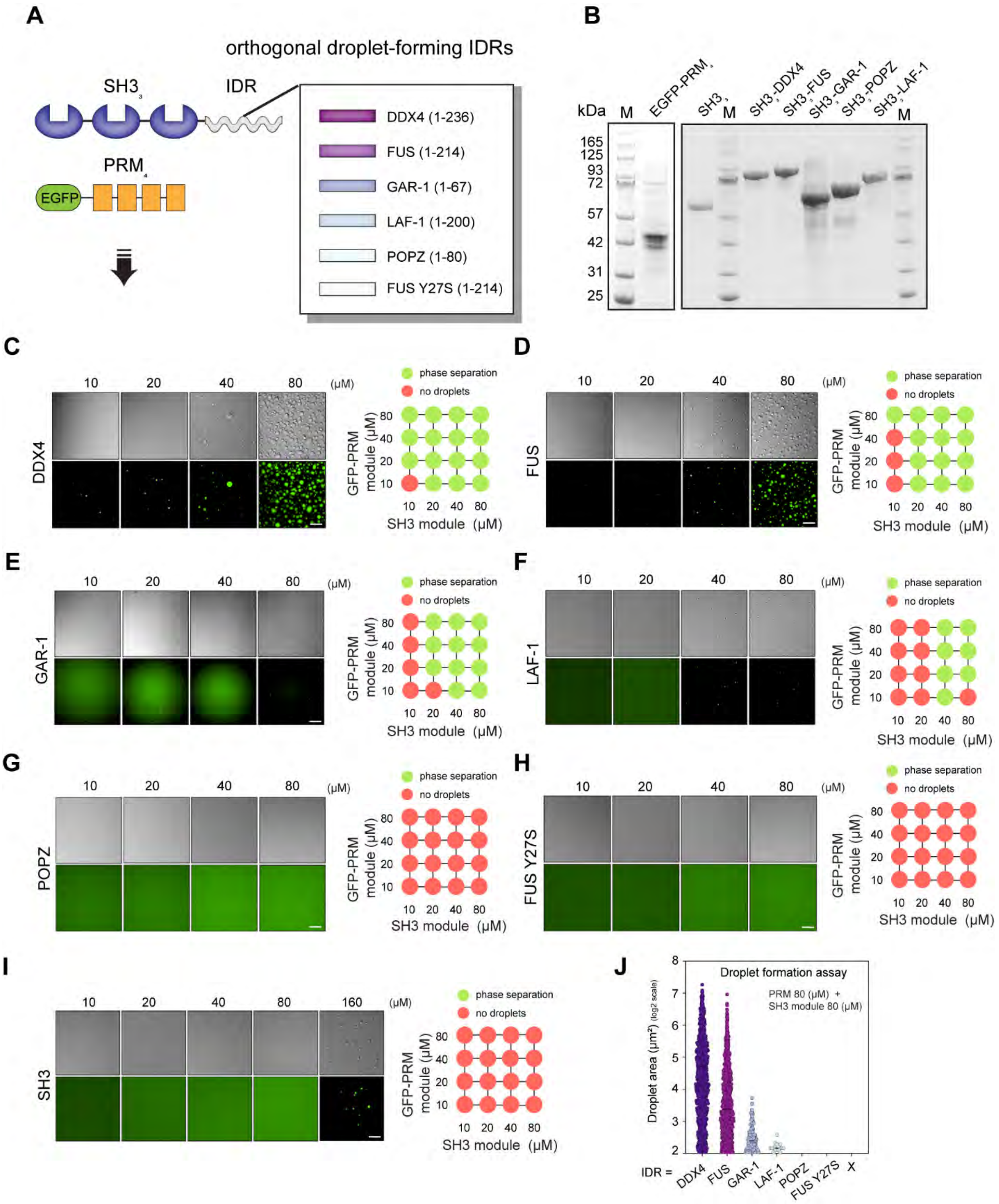


**Figure S2.** ***In vitro*** **droplet formation assays using the SH3–PRM multivalent interaction system fused to orthogonal intrinsically disordered regions (IDRs).**

(A) Schematic of $SH3_3$-IDR fusion proteins and EGFP-$PRM_4$ constructs used for phase separation assays. IDRs tested include DDX4 (1–236), FUS (1–214), GAR1 (1–67), LAF1 (1–200), POPZ (1–80), and the phase-separation-deficient mutant FUS Y27S (1–214).

(B) Coomassie-stained SDS-PAGE analysis of purified proteins used in this study.

(C–I) Representative brightfield and fluorescence microscopy images of in vitro droplet formation assays performed at the indicated SH3 module and GFP-PRM module concentrations. Green fluorescence corresponds to SH33-PRM4 condensates. In the phase diagrams, green circles indicate phase separation, whereas red circles indicate no detectable droplets. Scale bar: 20 μm. (I) Quantification of droplet area under conditions containing 80 μM SH3 module and 80 μM PRM module. Data are shown on a log2 scale.

**Represenative populations sampling D-IDR x E-WT RIPPLEs across different expression levels**

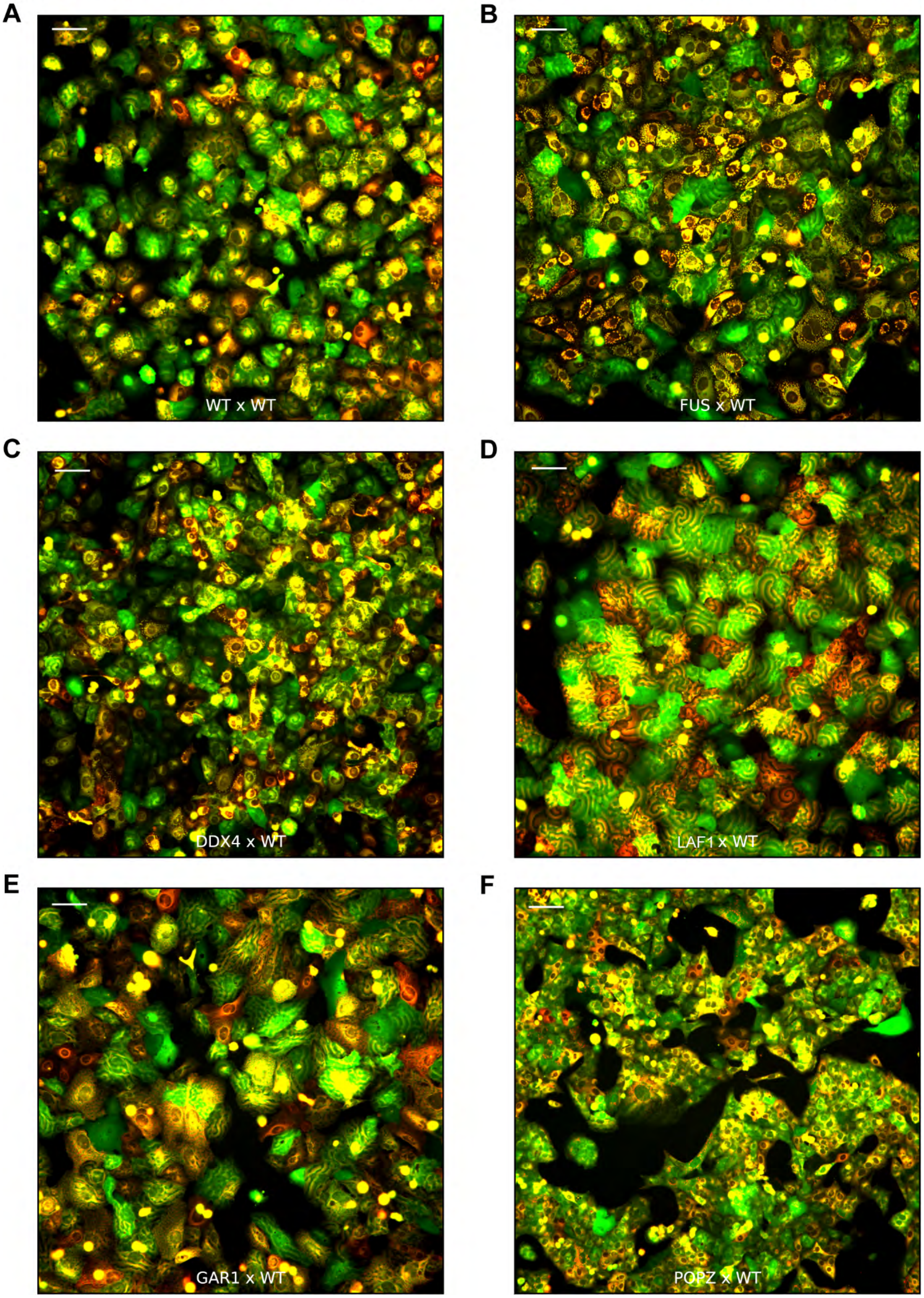

**Figure S3. Population sampling of IDR-MinD/MinE-WT RIPPLE designs.** Representative whole-field fluorescence images of IDR-MinD/MinE-WT RIPPLE populations sampled across different MinD/MinE expression levels. Scale bars, 100 µm. (A) WT-MinD/MinE-WT, (B) FUS-MinD/MinE-WT, (C) DDX4-MinD/MinE-WT, (D) LAF1-MinD/MinE-WT, (E) GAR1-MinD/MinE-WT, and (F) POPZ-MinD/MinE-WT.

**Represenative populations sampling D-WT x E-IDR RIPPLEs across different expression levels**

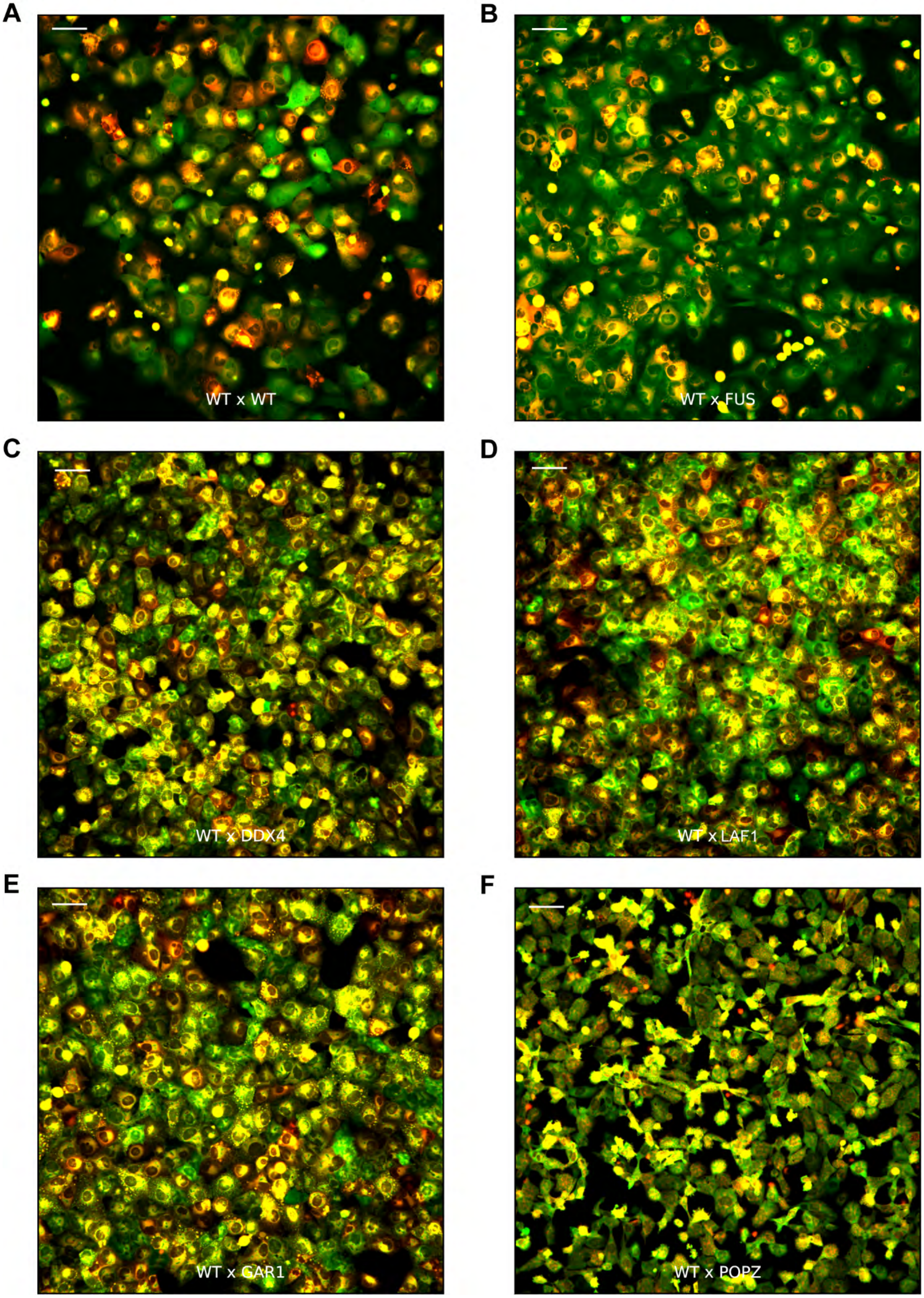

**Figure S4. Population sampling of WT-MinD/MinE-IDR RIPPLE designs.** Representative whole-field fluorescence images of WT-MinD/MinE-IDR RIPPLE populations sampled across different MinD/MinE expression levels. Scale bars, 100 µm. (A) WT-MinD/MinE-WT, (B) WT-MinD/MinE-FUS, (C) WT-MinD/MinE-DDX4, (D) WT-MinD/MinE-LAF1, (E) WT-MinD/MinE-GAR1, and (F) WT-MinD/MinE-POPZ.

Represenative populations sampling dual-IDR RIPPLE combinatorial library members across different expression levels

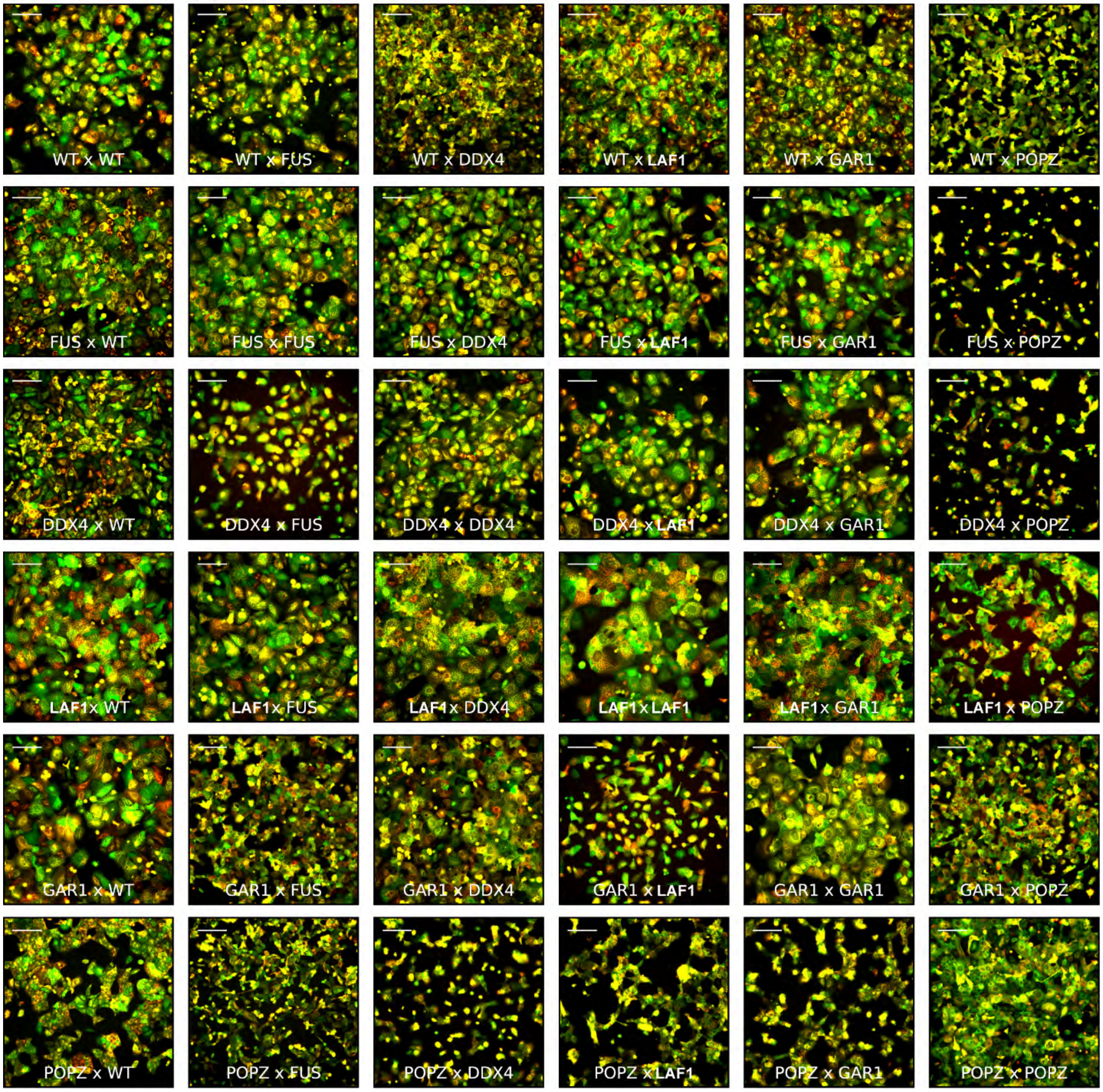


**Figure S5. Population sampling of dual-IDR RIPPLE designs.** Representative whole-field fluorescence images of dual-IDR (IDR-MinD/MinE-IDR) RIPPLE populations sampled across different MinD/MinE expression levels. Scale bars, 200 μm. Rows and columns correspond to the indicated IDR sequences in the following order: WT (non-IDR), FUS, DDX4, LAF1, GAR1, and POPZ.

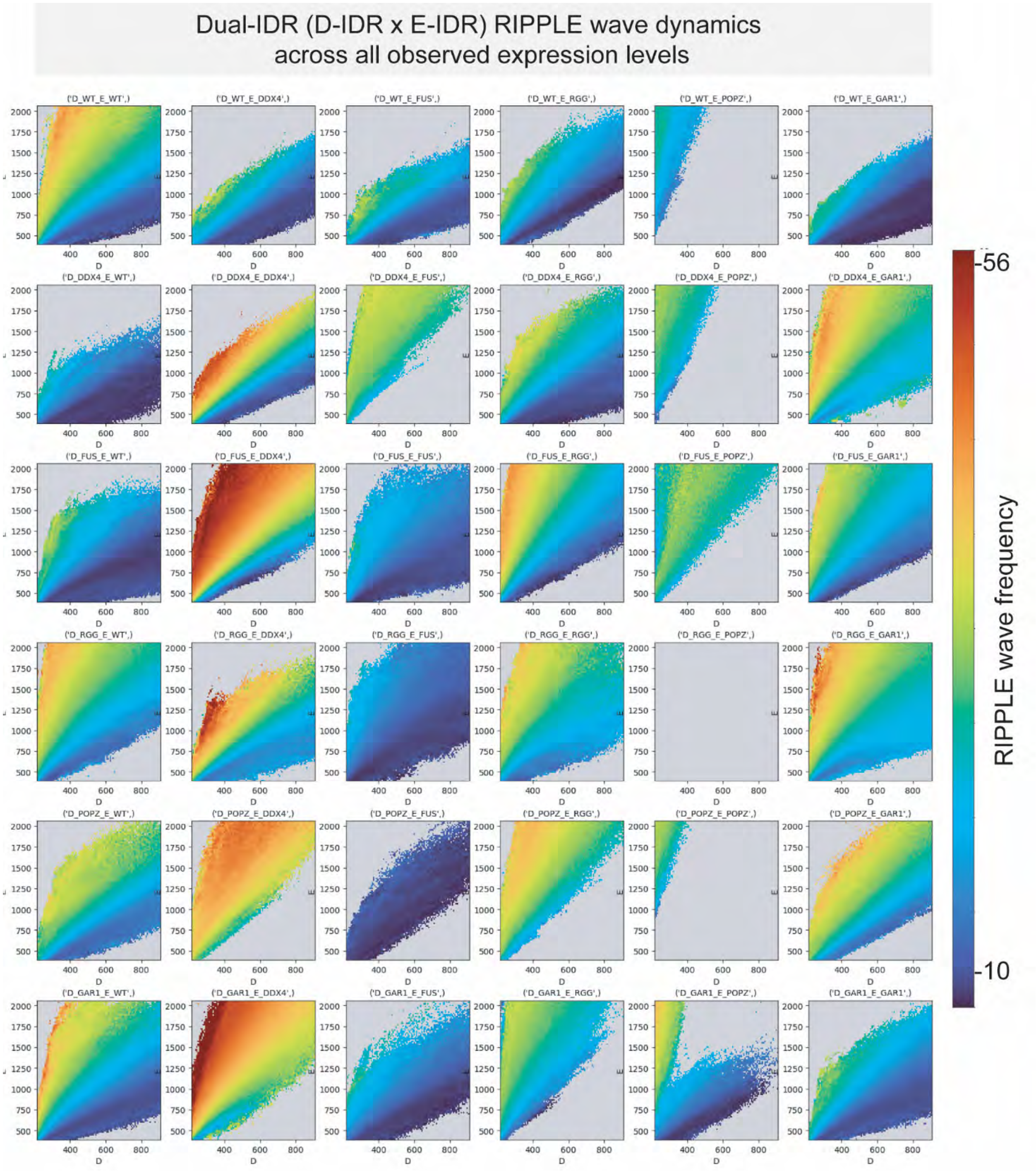


**Figure S6. Dual-IDR RIPPLE wave dynamics across population with all observed expression levels.** Distinct wave-dynamic phase portraits of dual-IDR (IDR-MinD/MinE-IDR) RIPPLE populations across all observed expression levels, shown as binned 2D histograms colored by mean oscillation frequency. Rows and columns correspond to the indicated IDR sequences in the following order: WT (non-IDR), DDX4, FUS, POPZ, GAR1, and LAF1.

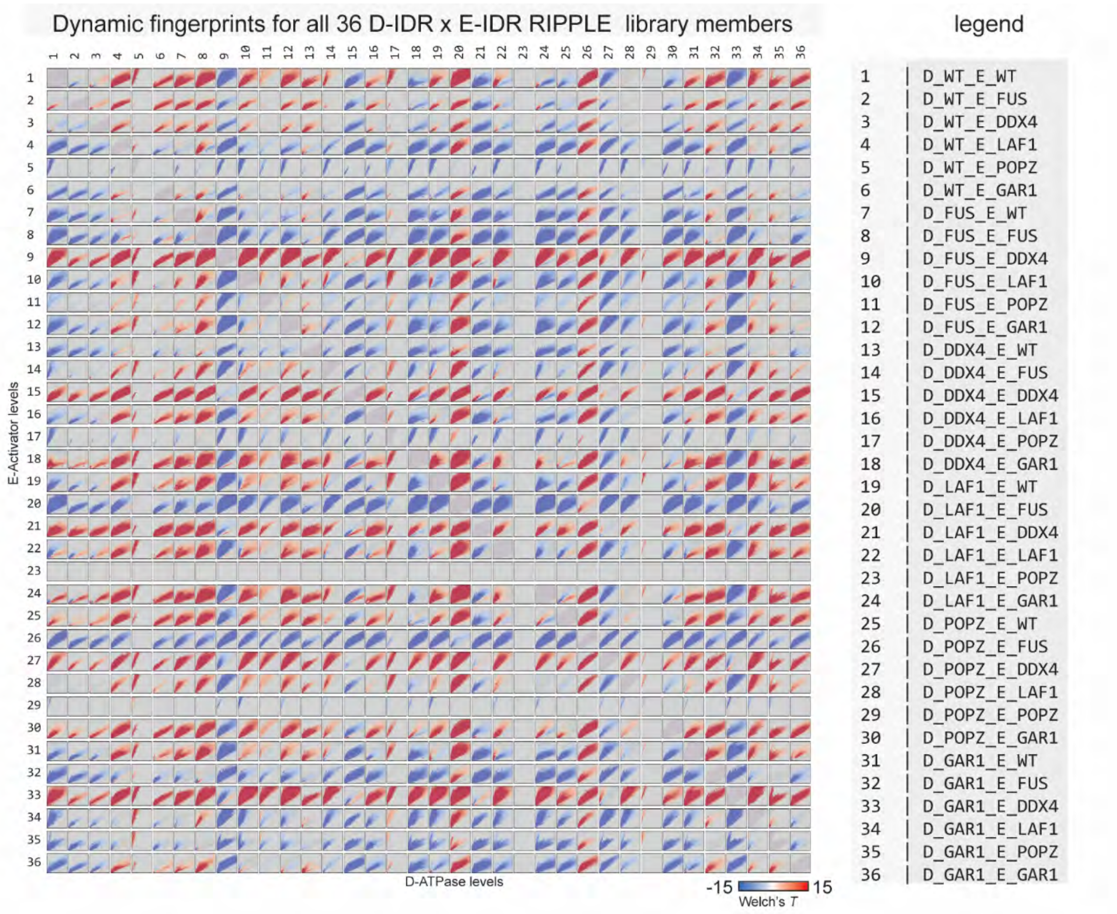
Dynamic fingerprints for all 36 D-IDR x E-IDR RIPPLE library members
legend
E-Activator levels
D-ATPase levels
-15
15
Welch's T
1 | D_WT_E_WT
2 | D_WT_E_FUS
3 | D_WT_E_DDX4
4 | D_WT_E_LAF1
5 | D_WT_E_POPZ
6 | D_WT_E_GAR1
7 | D_FUS_E_WT
8 | D_FUS_E_FUS
9 | D_FUS_E_DDX4
10 | D_FUS_E_LAF1
11 | D_FUS_E_POPZ
12 | D_FUS_E_GAR1
13 | D_DDX4_E_WT
14 | D_DDX4_E_FUS
15 | D_DDX4_E_DDX4
16 | D_DDX4_E_LAF1
17 | D_DDX4_E_POPZ
18 | D_DDX4_E_GAR1
19 | D_LAF1_E_WT
20 | D_LAF1_E_FUS
21 | D_LAF1_E_DDX4
22 | D_LAF1_E_LAF1
23 | D_LAF1_E_POPZ
24 | D_LAF1_E_GAR1
25 | D_POPZ_E_WT
26 | D_POPZ_E_FUS
27 | D_POPZ_E_DDX4
28 | D_POPZ_E_LAF1
29 | D_POPZ_E_POPZ
30 | D_POPZ_E_GAR1
31 | D_GAR1_E_WT
32 | D_GAR1_E_FUS
33 | D_GAR1_E_DDX4
34 | D_GAR1_E_LAF1
35 | D_GAR1_E_POPZ
36 | D_GAR1_E_GAR1

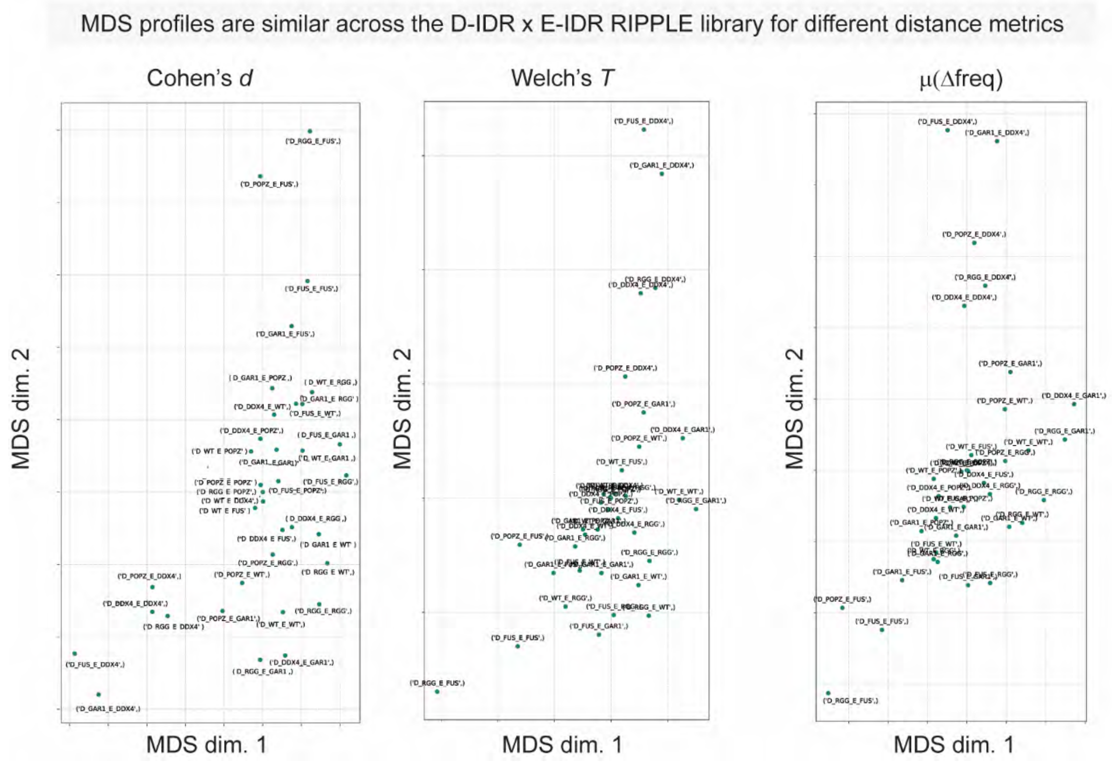
MDS profiles are similar across the D-IDR x E-IDR RIPPLE library for different distance metrics
Cohen's d
Welch's T
μ(Δfreq)
MDS dim. 2
MDS dim. 1

**Figure S7. Pairwise comparison of dual-IDR RIPPLE dynamic fingerprints.**

(A) Pairwise comparison of dynamic fingerprints from dual-IDR (IDR-MinD/MinE-IDR) RIPPLE designs across all observed expression levels. Dynamic fingerprints, represented as 2D histograms relating MinD and MinE expression levels to oscillation frequency, were compared on a bin-by-bin basis using Welch's t-statistic. Each histogram bin was recolored according to its corresponding t-statistic, where red indicates faster oscillations and blue indicates slower oscillations relative to the comparison design. Diagonal entries (self-comparisons) are shown in gray, and pairwise comparisons are symmetric across the diagonal with opposite statistic signs. Numerical labels correspond to dual-IDR designs indicated in the legend.

(B) Effect of distance metric choice on the organization of dual-IDR RIPPLE dynamic fingerprints in multidimensional scaling (MDS) space. Pairwise distances between dynamic fingerprints were calculated using Cohen's d, Welch's t-statistic, or absolute frequency difference:

$$d_{cohen} = \frac{\overline{X_1} - \overline{X_2}}{s_{pooled}} \qquad T_{Welch} = \frac{\overline{X_1} - \overline{X_2}}{\sqrt{\frac{s_1^2}{n_1} + \frac{s_2^2}{n_2}}} \qquad \Delta freq = \overline{f_1} - \overline{f_2}$$

Distances were squared and summed across all histogram bins to generate pairwise similarity matrices for MDS analysis. The resulting 3D embeddings are shown with the x- and y-axes corresponding to the first two MDS dimensions. All three distance metrics produced qualitatively similar trajectories in MDS space.

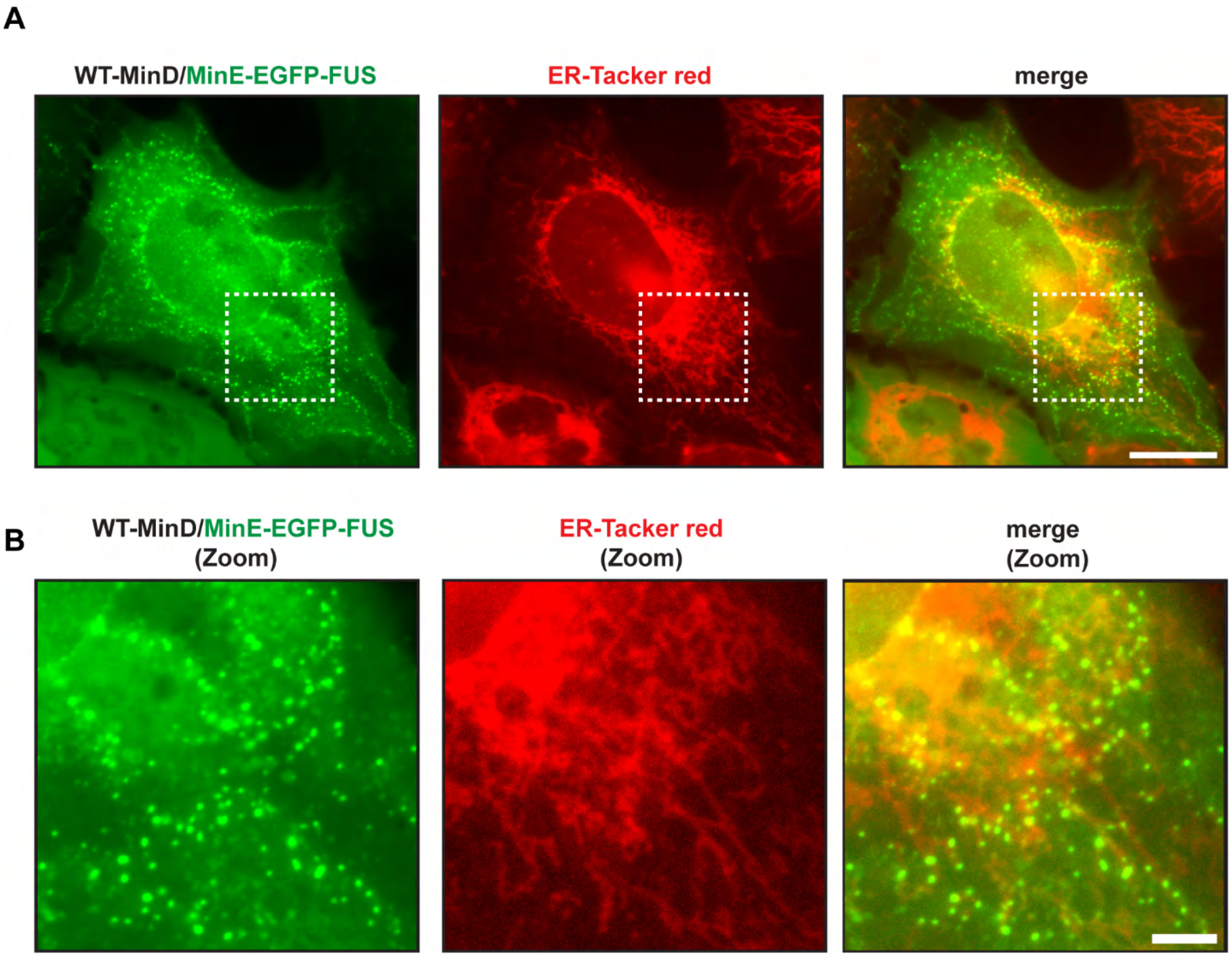


**Figure S8. Live cell Imaging of Activator-IDR RIPPLE and ER Tracker** Red Fluorescence images show the distribution of WT-MinD/MinE-EGFP-FUS-IDR alongside ER-Tracker Red staining to highlight the ER network. The merged image reveals co-localization of MinE-FUS-IDR active droplets with ER structures. Scale bar: 20 μm. The white dashed box indicates the region further magnified in the bottom row. Scale bar: 5 μm.

**Supplementary videos:**

**Movie S1:** Targeting FUS-IDR to MinD produces a condensed wavefront in a MinDE reaction-diffusion system.

**Movie S2:** Targeting FUS-IDR to the MinE-Activator produces droplets at the rear in the wake of a MinDE reaction-diffusion wave.

**Movie S3:** Representative D-IDR RIPPLE dynamics across diverse reaction-diffusion waveforms. Systematic analysis of D-IDR RIPPLEs across 1000s of reaction-diffusion driving waveforms defines frequency-dependent transitions between dilute and condensed states from different IDR sequences.

**Movie S4:** Representative E-IDR RIPPLE dynamics across diverse reaction-diffusion waveforms. Systematic analysis of E-IDR RIPPLEs across 1000s of reaction-diffusion driving waveforms defines dynamic bandpass patterning of droplet arrays, lattices, and phase-separated macrostructures.

**Movie S5:** Emulsification of highly condensed D-IDR RIPPLEs by co-expression of matched or unmatched E-IDR partners. Examples of emulsification of highly condensed D-IDR RIPPLEs by co-expression of a matched or unmatched E-IDR partner.

**Movie S6:** Dual-IDR RIPPLE combinations produce diverse persistent phase-separated macrostructures (GAR-1 panel).

**Movie S7:** High-magnification imaging of dual-IDR RIPPLE combinations producing dynamic emulsions (E-LAF1 panel).

**Movie S8:** High-magnification imaging of exotic patterning phenotypes and behaviors observed across the combinatorial dual-IDR RIPPLE library.

**Movie S9:** Recruitment of macromolecules to RIPPLE condensates, patterns, and macrostructures.
